# Development and utilization of *Treponema pallidum* expressing green fluorescent protein to study spirochete-host interactions and antibody-mediated clearance: expanding the toolbox for syphilis research

**DOI:** 10.1101/2024.10.21.619476

**Authors:** Kristina N. Delgado, Crystal F. Vicente, Christopher M. Hennelly, Farhang Aghakhanian, Jonathan B. Parr, Kevin P. Claffey, Justin D. Radolf, Kelly L. Hawley, Melissa J. Caimano

## Abstract

Syphilis is a sexually transmitted infection caused by the highly invasive and immunoevasive spirochetal pathogen *Treponema pallidum* subsp. *pallidum* (*TPA*). Untreated syphilis can lead to infection of multiple organ systems, including the central nervous system. The alarming increase in syphilis cases globally underscores the importance of developing novel strategies to understand the complexities of syphilis pathogenesis. In this study, we took advantage of recent advances in *in vitro* cultivation and genetic manipulation of syphilis spirochetes to engineer a *TPA* strain that constitutively expresses green fluorescent protein (GFP). GFP^+^ *TPA* grew identically to the Nichols parent strain *in vitro* and exhibited wild-type infectivity in the rabbit model. We then used the GFP^+^ strain to visualize *TPA* interactions with host cells during co-cultivation *in vitro*, within infected rabbit testes, and following opsonophagocytosis by murine bone marrow-derived macrophages. Development of fluorescent strain also enabled us to develop a flow cytometric-based assay to assess antibody-mediated damage to the spirochete’s fragile outer membrane (OM), demonstrating dose-dependent growth inhibition and OM disruption *in vitro*. Notably, we observed greater OM disruption of GFP^+^ *TPA* with sera from immune rabbits infected with the *TPA* Nichols strain compared to sera generated against the genetically distinct SS14 strain. These latter findings highlight the importance of OM protein-specific antibody responses for clearance of *TPA* during syphilitic infection. The availability of fluorescent *TPA* strains paves the way for future studies investigating spirochete-host interactions as well as functional characterization of antibodies directed treponemal OM proteins, the presumptive targets for protective immunity.

**Importance:** Syphilis, a sexually transmitted infection caused by *Treponema pallidum* (*TPA*), remains a pressing threat to global public health. *TPA* has a remarkable and still poorly understood ability to disseminate rapidly from the site of inoculation and establish persistent infection throughout the body. Recent advances in *in vitro* cultivation and genetic manipulation of syphilis spirochetes enabled the development of fluorescent *TPA*. In the study, we generated and characterized an infectious *TPA* strain that constitutively expresses green fluorescent protein and used this strain to visualize interaction of *TPA* with host cells and functionally characterize antibodies directed against treponemal outer membrane proteins. Most notably, we assessed the ability of surface-bound antibodies to inhibit growth of *TPA in vitro* and/or disrupt the spirochete’s fragile outer membrane. Fluorescent *TPA* strains provide a powerful new tool for elucidating host-pathogen interactions that enable the syphilis spirochete to establish infection and persistent long-term within its obligate human host.

## Introduction

Syphilis is a complex, multi-phase sexually transmitted disease caused by the highly invasive and immunoevasive spirochete *Treponema pallidum* subsp. *pallidum* (*TPA*) (1, 2). Following colonization of skin and mucosal surfaces, early and widespread hematogenous dissemination of spirochetes is the rule during acquired human syphilis, as evidenced by the many organ systems, including the central nervous system, *TPA* invades to establish persistent, often lifelong, infection (1, 2). No form of the disease better exemplifies *TPA*’s invasiveness than gestational syphilis (2, 3). When syphilis is acquired during pregnancy, *TPA* readily penetrates the fetal-placental barrier, often giving rise to serious consequences in the unborn offspring, including demise of the fetus or neonate (2, 4). Despite efforts by the Centers for Disease Control and Prevention (CDC) and World Health Organization to curtail the spread of syphilis, its incidence continues to increase in the United States and worldwide (1, 2, 5, 6). As of 2022, the last year for which complete data are available from the CDC, more than 200,000 cases of syphilis were reported in the United States, representing a ∼17% increase compared to 2021 (7). The same year also saw 3,761 reported cases of congenital syphilis, a ∼32% increase from 2021 (8). These alarming trends underscore the importance of developing novel approaches to better understand the complexities of syphilis pathogenesis and the mechanisms underlying protective immune responses (1, 2).

The historical inability to propagate *TPA* continuously *in vitro* has been a major roadblock for developing genetic tools to identify *TPA* virulence determinants and characterize host-pathogen interactions *in vitro* and in the experimental rabbit model (9, 10). In 1981, Fieldsteel, Cox, and Moeckli (11) reported that coculture of *TPA* with Sf1Ep cottontail rabbit skin epithelial cells in modified tissue culture media under microaerophilic conditions improved survival of *TPA in vitro*, supporting up to 100-fold multiplication, but could not sustain growth long-term. Subsequent refinement of this coculture system by Edmondson, Norris, and colleagues (12) in 2018 enabled reproducible, long-term replication of *TPA in vitro*. This groundbreaking accomplishment was followed not long after by the first reported genetic manipulation of *in vitro*-cultivated *TPA* by chemical transformation (13, 14). Bacteria genetically modified to express fluorescent reporters have been instrumental for tracking pathogens, including other spirochetes, as they bind to and penetrate diverse cell types and tissues *in vitro* and *in vivo* (15–20). Grillova *et al*. (21) recently described a *TPA* strain SS14 expressing red-shifted green fluorescent protein, establishing the feasibility of using fluorescent reporters for syphilis research. Herein, we build upon this pioneering report by generating a green fluorescent strain in the genetically distinct *TPA* Nichols background. We then used this GFP^+^ strain to visualize host-pathogen interactions during co-cultivation *in vitro*, within infected rabbit testes, and following opsonophagocytosis by murine bone marrow-derived macrophages. We also developed a flow cytometry-based assay to assess antibody-mediated growth inhibition and/or disruption of the syphilis spirochete’s fragile outer membrane (OM), presumptive markers for bacterial killing in the rabbit model and, presumably, humans. Our findings illustrate how fluorescently labelled *TPA* strains can be used to investigate host-pathogen interactions and clearance of treponemes during syphilitic infection.

## RESULTS

### *TPA* constitutively expressing green fluorescent protein exhibits wild-type viability and cellular adhesiveness

To genetically engineer a green fluorescent *TPA* strain, we used the strategy described by Romeis *et al.* (13) to transform the WT *TPA* Nichols strain with a suicide vector encoding tandem Extra-superfolder green fluorescent protein (GFP) (22) and kanamycin-resistance (*kanR*) transgenes in place of the native *tprA locus* (pMC5836; Fig. S1A). Extra-superfolder GFP was selected for these studies as it folds more efficiently and fluoresces brighter than enhanced GFP (23, 24). As a control, we also transformed WT *TPA* Nichols with a version of the suicide vector encoding only the *kanR* transgene (pMC5722; Fig. S1B). Following recovery, transformants were passaged in TpCM-2 medium containing 200 μg/ml kanamycin (TpCM-2Kan). By the third passage *in vitro* (day 42), motile treponemes were observed for *TPA* transformed with *kanR* and *gfp-kanR* constructs but only the latter were fluorescent (GFP^+^) (Fig. 1A and Video S1-S2). Replacement of *tprA* in both strains was confirmed by PCR amplification of *TPA* genomic DNA (Fig. S1C-D and Table S1). As shown in Fig. 1B, WT, *kanR* and GFP^+^ *TPA* displayed highly similar growth profiles over 12 passages (168 days). Consistent with prior studies using nonfluorescent *TPA* (25), confocal composite images (Fig. 1C-D and Video S3) of *in vitro*-cultivated GFP^+^ *TPA* showed numerous treponemes on the surfaces of Sf1Ep cells; despite exhaustive efforts, we were unable to find any intracellular organisms.

**Fig 1.**
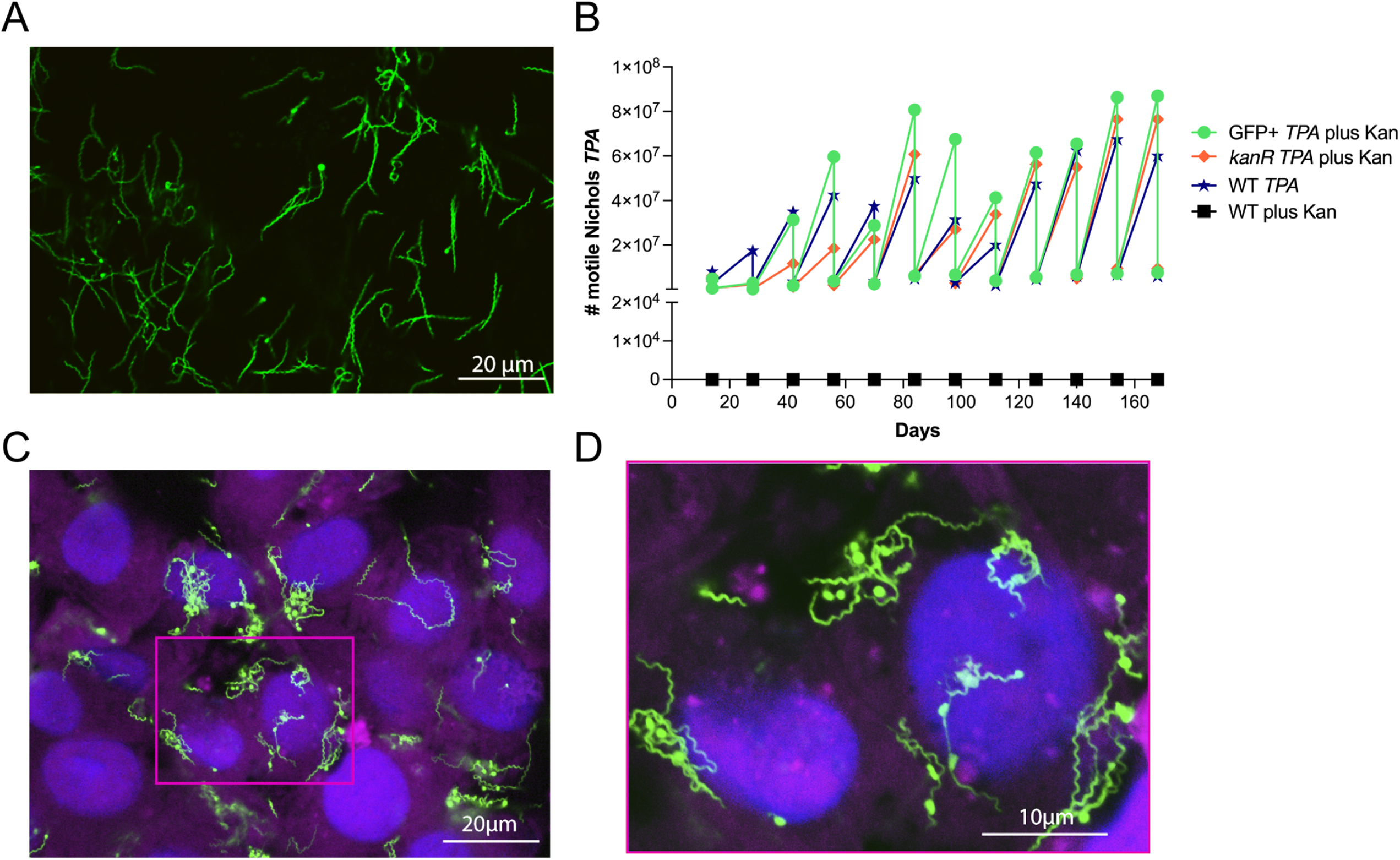
GFP^+^ *TPA* Nichols grows comparably to WT *in vitro* and localizes predominantly to the surface of rabbit epithelial cells. (**A**) Representative epifluorescence image of *in vitro*-cultivated GFP^+^ *TPA* in suspension following harvest from Sf1Ep cells using trypsin-EDTA. (**B**) Growth curves for GFP^+^ (green) and *kanR* (orange) *TPA* in the presence of kanamycin and the wild-type (WT) parent grown in the presence (black) and absence (blue) of kanamycin. (**C**) Representative confocal image (1 μm optical section) of GFP^+^ *TPA* co-cultured with Sf1Ep cells labeled with DAPI (blue) and Cholera Toxin AF647 (magenta), which stain host cell nuclei and plasma membranes, respectively. (**D**) Digital enlargement (3X) of boxed area in **C**. Z-stack of individual 1 μm optical sections showing surface localization of *TPA* can be found in Video S3.

To evaluate expression of GFP in *TPA* harvested from Sf1Ep cells, we employed flow cytometry, a technique used extensively by us (19, 20, 26) and others (27–29) to track fluorescent *Borrelia burgdorferi*, the Lyme disease spirochete. Given the narrow width (∼0.2 nm) (30) and elongated, helical waveform morphology of *TPA* (12, 31), separating spirochete populations from background noise by light scattering proved to be challenging. Instead, we used staining of GFP^+^ *TPA* with propidium iodide (PI), a membrane-impermeant, fluorescent DNA dye, to exclude flow cytometric events caused by debris. After harvest, treponemes were fixed with 2% paraformaldehyde, permeabilized with 0.01% Triton X-100, and counterstained with PI. Using unstained WT *TPA* to set gating parameters, we eliminated GFP/PI double-negative events using a dump gate (Fig. S2A) and then established a gating strategy for PI^+^ and GFP^+^ using detergent-treated WT *TPA* stained with PI (Fig. S2B) and untreated GFP^+^ *TPA* (Fig. S2C). In the absence of detergent, <1% of GFP^+^ *TPA* stained with PI, confirming that trypsinization to release treponemes during harvest does not disrupt the spirochete’s delicate OM (32). After eliminating double-negative events, ∼97% of detergent-treated treponemes harvested from Sf1Ep cells were PI^+^ (Fig. 2A); ∼96% of these were GFP^+^ with a mean fluorescence intensity (MFI) of 7381± 346. Essentially identical results were obtained by gating first on GFP; 97.8% of GFP^+^ events also were PI^+^ (Fig. 2B). Collectively, these data confirm the feasibility of using flow cytometry to quantitate fluorescent *TPA* populations.

**Fig 2.**
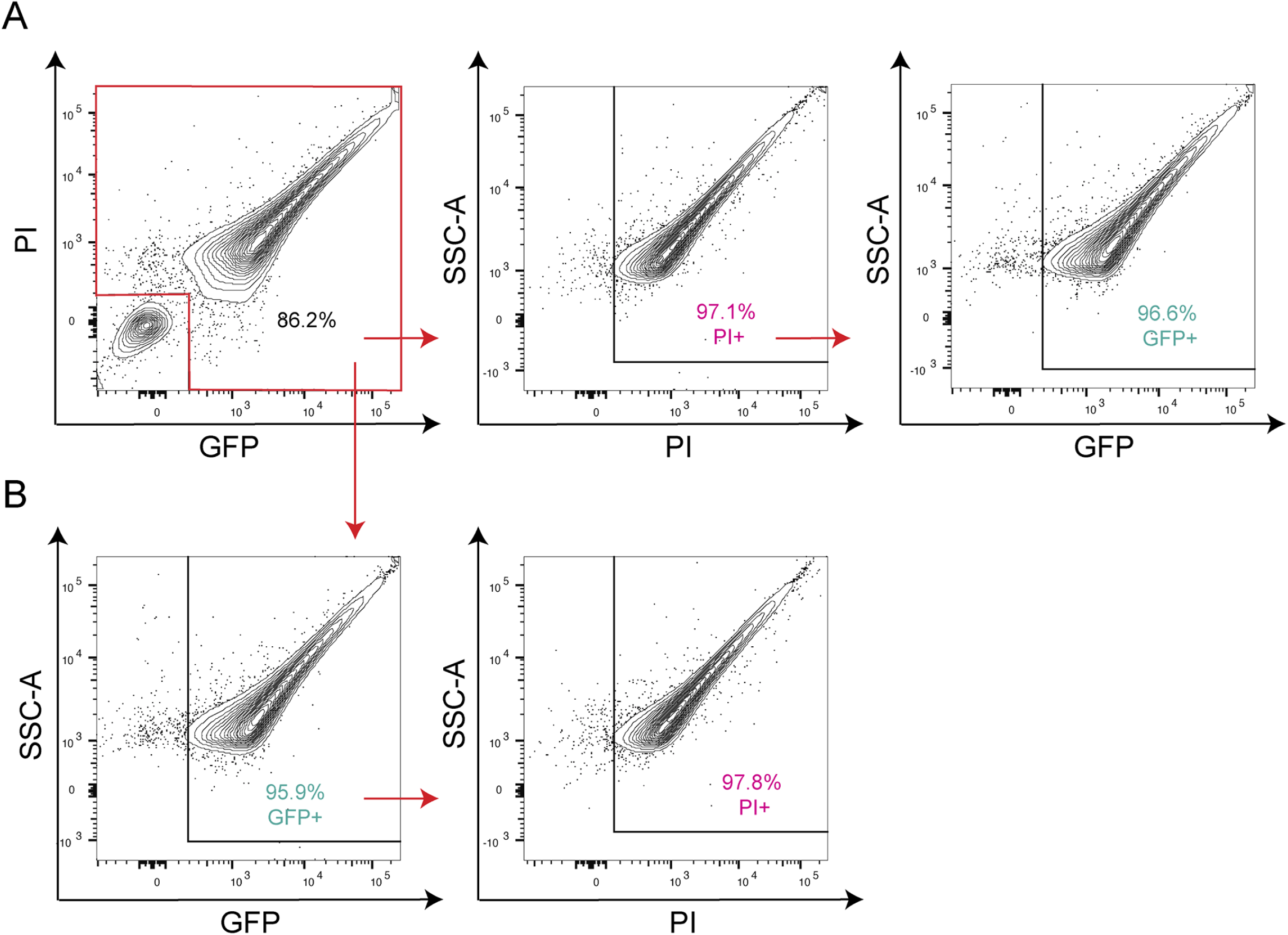
Flow cytometric analysis of GFP expression by GFP^+^ *TPA* during *in vitro* cultivation. Flow cytometry of GFP^+^ *TPA* dissociated from Sf1Ep cells using Trypsin-EDTA (see Methods). Gating strategies used to eliminate double-negative background events and define PI^+^ and GFP^+^ populations are shown in Fig. S2. Events within the red boxed area in the top left panel were used to quantify (**A**) GFP^+^ events within the PI^+^ population and (**B**) PI^+^ events within the GFP^+^ population. Results are representative of three independent experiments.

### Whole-genome sequencing to confirm replacement of *tprA* with *gfp*-*kanR* in *TPA* Nichols

Whole-genome sequencing (WGS) was performed on DNA extracted from *in vitro*-cultivated GFP^+^ *TPA* at passages 6, 9 and 12 to confirm replacement of *tprA* with the *gfp*-*kanR* cassette and rule out additional insertions of the cassette elsewhere in the genome. Using the Nichols reference genome (CP004010.2), few if any reads mapped to the *tprA* locus in the GFP^+^ *TPA* strain (Fig. S3). In contrast, abundant reads mapped to this region using a Nichols genome modified *in silico* to contain the *gfp*-*kanR* cassette (Fig. S3). Except for reads mapping to the *tpp47* and *flaA1* promoters used to drive transcription of *kanR* and *gfp*, respectively, we saw no evidence for insertion of either transgene elsewhere in the chromosome. Searches for the pUC19 backbone used to generate pMC5836 yielded only spurious reads. A comparison of WGS data for GFP^+^ *TPA* passages 6, 9 and 12 detected a total of 28 polymorphisms in at least one *in vitro* passage compared to the *TPA* Nichols NCBI reference genome. Fifteen were present in the parental strain (*TPA* Nichols-Farmington) used to generate the GFP^+^ (Table S2). Of the remaining 13 differences, all but one were present in all three passages. Most of the differences in the GFP^+^ strain were either single nucleotide polymorphisms (SNPs) or small (1- to 2-bp) indels within homopolymeric regions that result in a frameshift mutation. Only one frameshift occurred within a gene (*tp0040/mcp1*) encoding a protein of known function. Of note, three of the frameshift mutations identified in the GFP^+^ strain but not the WT parent were identified independently by Edmondson *et al.* (33) in their *TPA* Nichols isolate. Importantly, no SNPs were detected in the *gfp*-*kanR* cassette in any of the passages. These data suggest that although the parental Nichols isolate used to generate the GFP^+^ strain is most likely non-clonal, the engineered strain is highly stable *in vitro*.

### GFP^+^ *TPA* exhibits wild-type infectivity in the rabbit model and comparable expression of GFP *in vitro* and following rabbit passage

To confirm that GFP^+^ *TPA* retained WT infectivity, we inoculated rabbits intratesticularly with a total of 2 × 10^7^ *in vitro-*cultivated GFP^+^ or WT *TPA*. In three independent experiments, rabbits inoculated with either strain developed orchitis within 12-14 days (Fig. 3A). Moreover, we saw no significant difference in the number of motile GFP^+^ and WT treponemes recovered at the time of harvest (*in vitro* → Rabbit 1; Fig. 3A and Video S4-5). Serial passage of each strain into a second rabbit yielded highly similar results with comparable values for both days to onset of orchitis and treponeme recovery (Rabbit 1 → Rabbit 2; Fig. 3A) in all three experiments. WGS analysis of GFP^+^ *TPA* recovered from rabbit testes confirmed that the *gfp-kanR* insertion was stable *in vivo* (Fig. S3), with only one additional SNPs, a 2-bp insertion within a non-coding region, were identified between *in vitro* passage 6 and rabbit passage 2 (Table S2); the single unique SNP, a 2-bp insertion in a polyG tract in the genome of Rabbit 1, occurred within a non-coding region. Confocal imaging of cryosections of testes obtained at the time of sacrifice revealed numerous GFP^+^ treponemes attached to the surface of testicular cells (Fig. 3B and Video S6); as in the *in vitro* studies, intracellular organisms were not identified in testes tissue. Lymph nodes and blood samples collected at the time of sacrifice from WT and GFP^+^ *TPA*-infected Rabbit 2 demonstrated comparable spirochete burdens by qPCR, confirming comparable hematogenous dissemination and invasion of distal tissues by the GFP^+^ strain (Fig. 3C).

**Fig 3.**
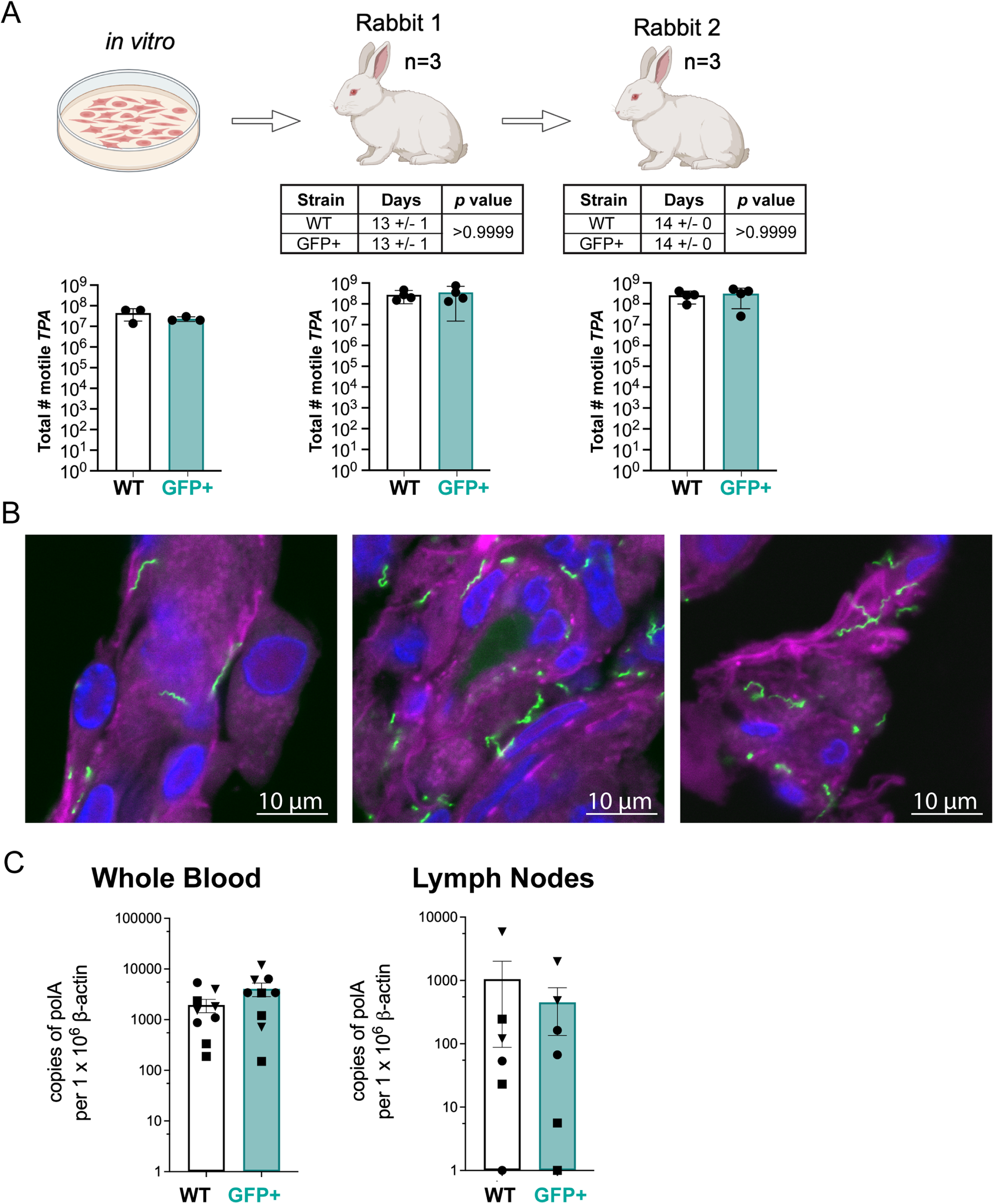
GFP^+^ *TPA* exhibits WT intratesticular growth, hematogenous dissemination, and localization to lymph nodes following intratesticular inoculation. (**A**) Schematic representation and results for serial passage of *in vitro*-cultivated WT (white) and GFP^+^ (cyan) *TPA* in rabbit testes. The total number of motile treponemes for each strain/condition were determined by darkfield microscopy. ‘Days’ indicates the times required for *TPA*-infected testes to reach peak orchitis; bars indicate the means ± standard deviations. No significant differences (*p* ≤ 0.05) were observed between WT and GFP^+^ *TPA* for either timing of orchitis or burdens in testes. Table and graphs represent data from three independent serial passage experiments. (**B**) Representative 1 μm optical sections of GFP^+^ *TPA* (green) revealing numerous extracellular treponemes in rabbit testes harvested at peak orchitis and stained with DAPI (blue) and Cholera Toxin Subunit B (magenta). Z-stacks of individual 1 μm optical sections showing surface localization of GFP^+^ *TPA* can be found in Video S6. (**C**) Bar graphs depicting spirochete burdens for WT (white) and GFP^+^ (cyan) *TPA* in blood (n = 3 replicates per rabbit, per strain, per experiment) and popliteal lymph nodes (n = 2 replicates per rabbit, per strain, per experiment) collected at sacrifice. Bars represent the means ± SEMs for *TPA polA* determined by qPCR normalized per 1 × 10^6^ copies of rabbit β-actin for Rabbit 2 for each strain from three independent experiments. Symbols (triangle, square and circle) represent data points from individual animals.

To evaluate whether the levels of GFP *in vivo* are comparable to those observed *in vitro*, we performed flow cytometry on treponemes from infected testes following harvest applying the same gating strategy described above. After removing double negative events, ∼95% of PI^+^ events were GFP^+^ with a mean MFI of 7583 ± 309 (Fig. 4A), essentially identical to the value (7381 ± 346, per above) following *in vitro* cultivation. Conversely, ∼95% of GFP^+^ events were PI^+^ when gating on the GFP^+^ population (Fig. 4B).

**Fig 4.**
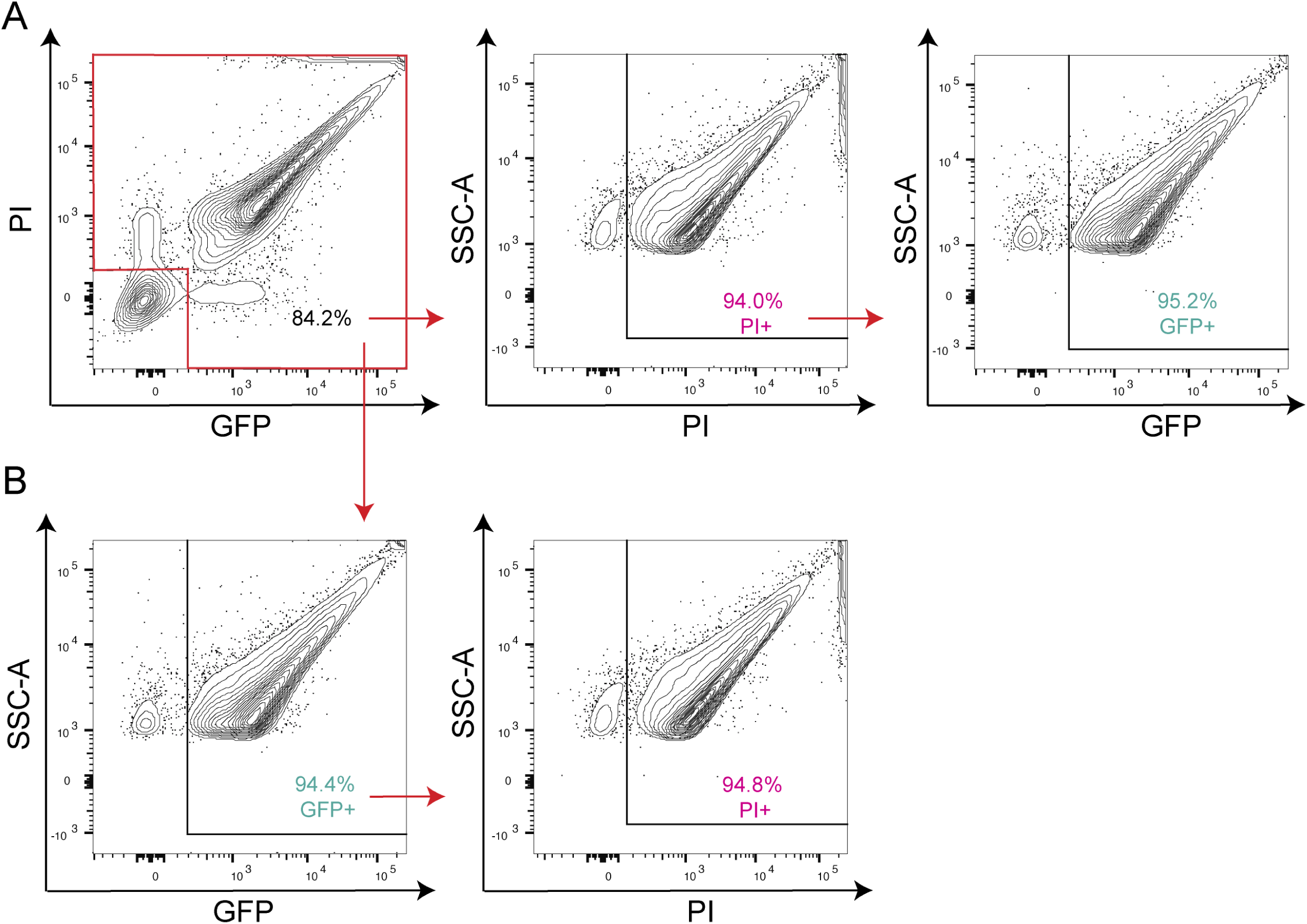
Expression of GFP by *TPA* harvested from rabbit testes. Flow cytometry of GFP^+^ *TPA* harvested from rabbit testes (*in vivo*), permeabilized with 0.01% Triton X-100, and counterstained with PI. Gating strategies used to eliminate non-spirochetal double-negative events and define PI^+^ and GFP^+^ populations are shown in Fig. S2. Events within the red boxed area in the top left panel were used to quantify (**A**) GFP^+^ events within the PI^+^ population and (**B**) PI^+^ events within the GFP^+^ population. Results are representative of three independent experiments.

Lastly, we assessed the ability of GFP^+^ *TPA* to cause cutaneous lesions following intradermal inoculation of rabbits (n = 3) with graded doses (10^5^ – 10^1^ per site) of both strains on either side of the same animal. Beginning 7 days post-inoculation (p.i.) until sacrifice on day 30, lesions were measured daily. As shown in Fig. 5A-B and Fig. S4, the overall size and time course for lesions elicited by WT and GFP^+^ *TPA* were not significantly different (*p* > 0.05). However, the lesions produced by GFP^+^ *TPA* at the 10^5^ dose were significantly (*p* ≤ 0.05) larger than those produced by the WT parent between days 14-23 p.i. (Fig. 5B). Lesions produced by WT and GFP^+^ *TPA* at the 10^4^-10^2^ doses were not significantly different over the course of the experiment (day 7 - day 30 p.i.). In contrast, lesions produced by WT *TPA* at the 10^1^ dose day 28-30 p.i. were larger than those for the GFP^+^ strain, possibly indicating a slight decrease in infectivity for the fluorescent strain. At the time of sacrifice (day 30 p.i.), we saw no significant differences in spirochete burdens based on normalized copy numbers of *polA* for the WT and GFP^+^ strains at sites inoculated with 10^4^ or 10^5^ organisms (Fig. 5C).

**Fig 5.**
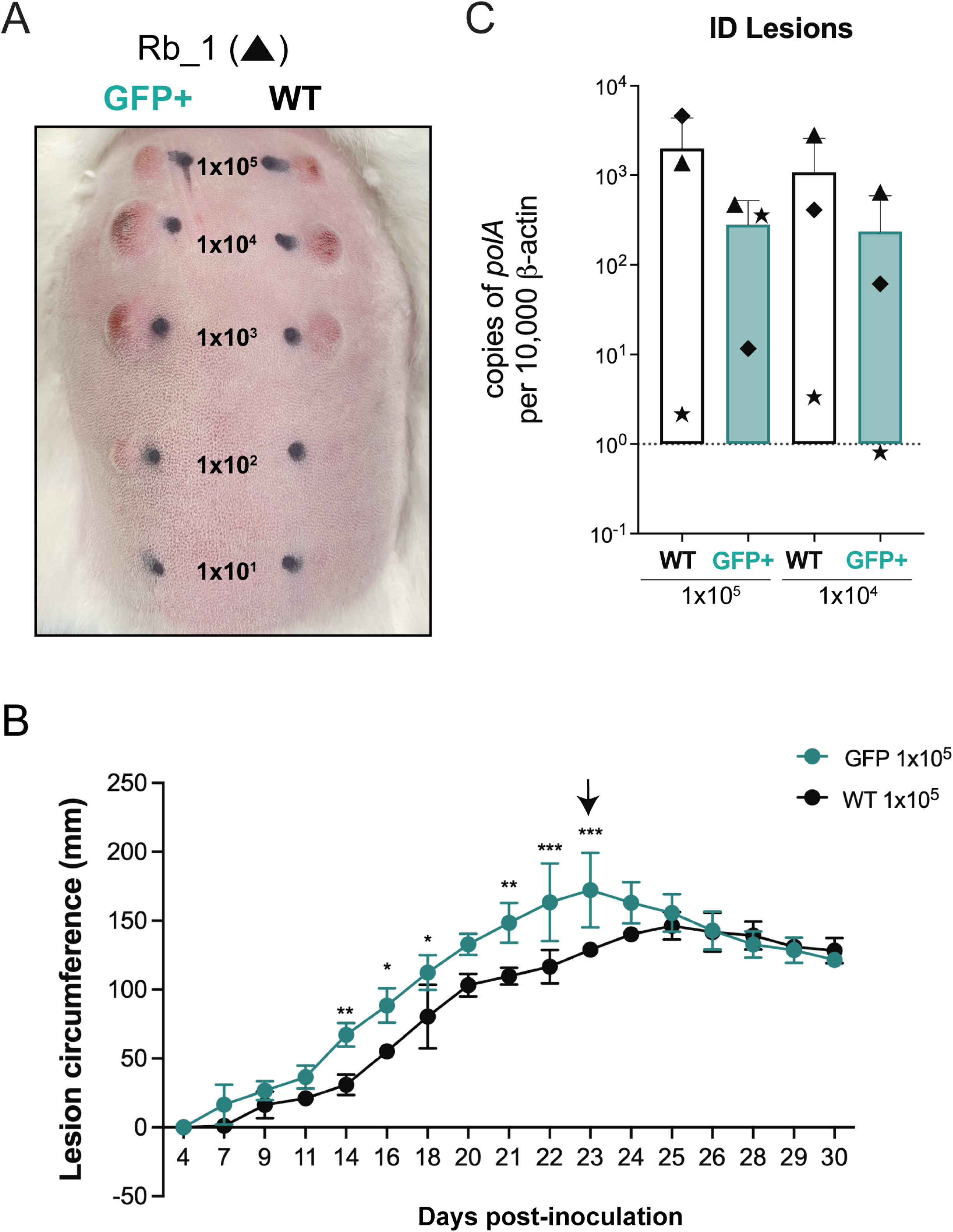
GFP^+^ *TPA* exhibits wild-type infectivity in rabbits by intradermal challenge. (**A**) Representative images of dermal lesions 23 days after intradermal inoculation with graded doses (1 × 10^5^ – 1 × 10^1^) of GFP^+^ (left) and WT (right) *TPA*. (**B**) Lesion circumferences (mm) for sites inoculated with 1 x 10^5^ GFP^+^ or WT *TPA*. Values represent the means ± standard deviations for three animals. *, *p* ≤ 0.05; **, *p* ≤ 0.01; ***, *p* ≤ 0.001. Arrow indicates the time point at which images in panel A were obtained. (**C**) Treponemal burdens in sites inoculated with 1 × 10^5^ and 1 × 10^4^ GFP^+^ or WT *TPA*. Bars represent the means ± standard deviations for *TPA polA* values normalized per 10^4^ copies of rabbit β-actin determined by qPCR. Symbols (triangle, square and circle) represent data points from three individual animals.

### Opsonophagocytosis of GFP^+^ *TPA* by murine bone marrow-derived macrophages

Macrophage-mediated opsonophagocytosis of *TPA* is widely considered to be critical for spirochete clearance (10). From the standpoint of vaccine development, opsonophagocytosis assays are essential for evaluating the potential protective capacity of antibodies directed against surface-exposed epitopes (34–36). We reasoned that incorporation of GFP^+^ *TPA* into this assay would expedite sample processing and analysis by eliminating the need to visualize macrophage-associated treponemes via indirect immunofluorescence. Accordingly, we used our recently developed *ex vivo* assay employing murine bone marrow-derived macrophages (37) to evaluate uptake and degradation of *in vitro*-cultivated GFP^+^ *TPA* following incubation with mouse syphilitic sera (MSS) generated against *TPA* Nichols or antiserum against the *TPA* Nichols BamA β-barrel extracellular loop 4 (α-BamA ECL4), both of which are strongly opsonic (37, 38). To preserve proteinaceous surface epitopes, treponemes used for opsonophagocytosis assay were recovered from Sf1Ep cells using Dissociation buffer rather than trypsin-EDTA (39). Normal mouse sera (NMS) and antisera against TP0751, a lipoprotein which, in our hands, is not an opsonic target (37, 40), were used as negative controls. Consistent with prior studies using nonfluorescent WT *TPA* harvested from rabbit testes (37, 38), preincubation of *in vitro*-cultivated GFP^+^ *TPA* with 10% MSS or α-BamA-ECL4 substantially increased internalization by BMDMs compared to NMS and α-TP0751 controls (Fig. 6A). For each serum, we determined the phagocytic index, a measure of opsonophagocytosis that considers the number of macrophages with ingested organisms as well as the number of treponemes phagocytosed per cell (37, 41). As shown in Fig. 6B, phagocytic indices for MSS and α-BamA ECL4 were significantly greater than those for NMS and α-TP0751, which were equivalent to the ‘no sera’ control.

**Fig 6.**
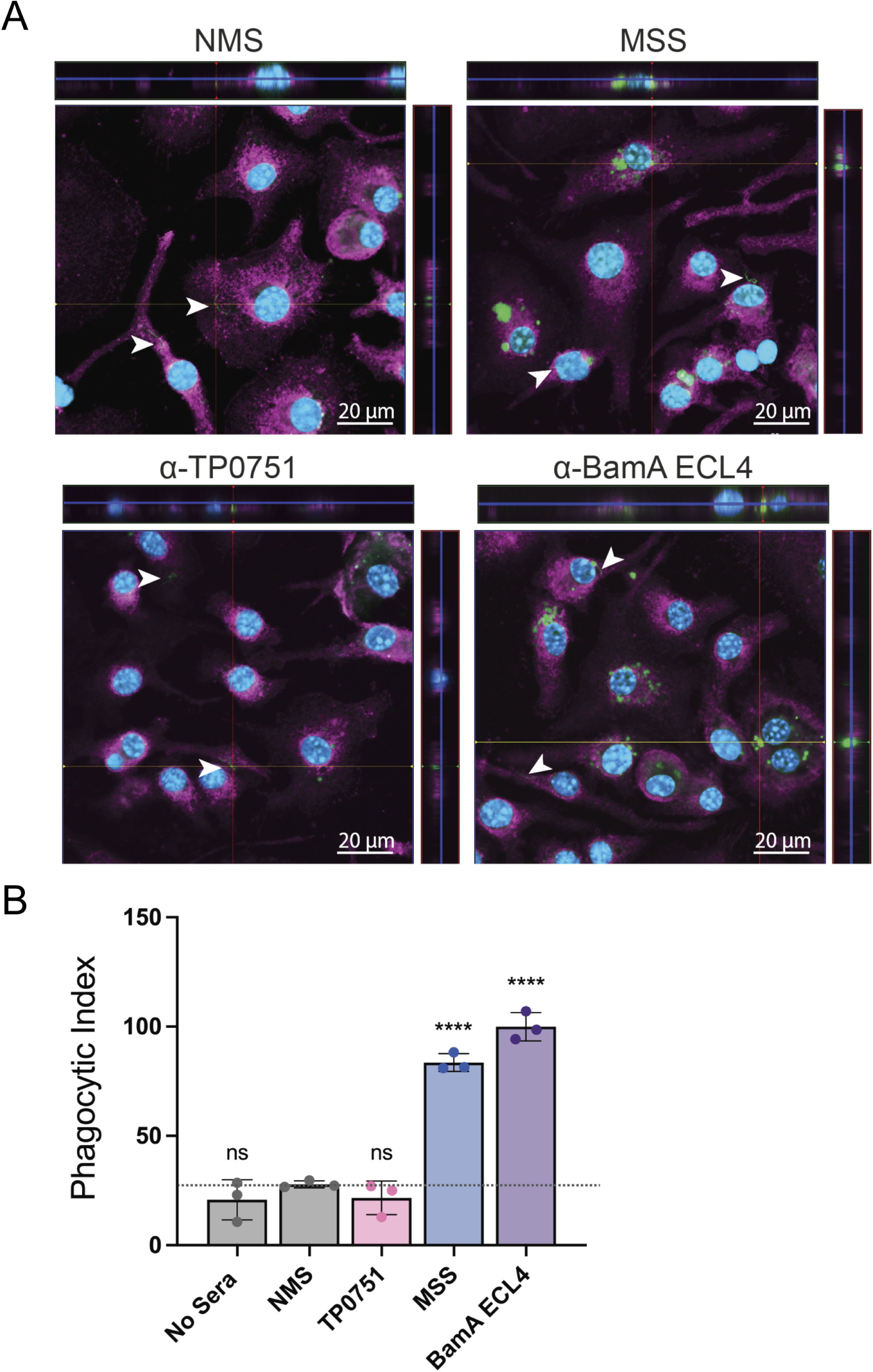
Opsonophagocytosis of GFP^+^ *TPA* by murine bone marrow-derived macrophages. (**A**) Representative 9-12 µm composite confocal images showing internalization and degradation of GFP^+^ *TPA (*green) by murine bone marrow-derived macrophages following pre-incubation with murine syphilitic serum (MSS) generated against *TPA* Nichols or mouse polyclonal antisera against BamA ECL4. Normal mouse sera (NMS) and mouse polyclonal antisera against TP0751, a non-opsonic periplasmic lipoprotein, were used as negative controls. White arrowheads indicate *TPA* on the surfaces of macrophages. Magenta, cholera toxin AF647; blue, DAPI. (**B**) Phagocytic indices for samples shown in panel A. Bars represent the mean ± standard deviation for three biological replicates per condition. ****, *p* ≤ 0.0001 compared to NMS.

### Immune rabbit serum and antibodies targeting BamA extracellular loop 4 inhibit growth and damage *TPA* OMs during *in vitro* cultivation

We recently demonstrated that sera from immune rabbits and antibodies targeting ECLs of specific *TPA* OM embedded β-barrel proteins (OMPs) can inhibit growth and even kill *TPA* Nichols in the absence of complement during *in vitro* cultivation (41). We, therefore, sought to determine if GFP^+^ *TPA* could be used to quantify the growth-inhibiting and/or bactericidal effects of α-ECL antibodies *in vitro* as an Fc receptor- and complement-independent ‘surrogate of protection’ for vaccine development. Consistent with our prior studies (41), neither NRS nor α-TP0751 antibodies had a measurable impact on growth *in vitro* (Fig. 7A). In contrast, IRS generated using the Nichols strain and antisera directed against ECL4 of the Nichols BamA dramatically (*p* ≤ 0.0001) reduced spirochete numbers to levels at or below the starting inoculum (Fig. 7A). We reasoned that damage to *TPA*’s fragile OM (35) during incubation with surface-directed antibodies could be used as a marker for bacterial killing. To investigate how co-cultivation with different antibodies affects OM integrity, we took advantage of our finding that intact GFP^+^ *TPA* exclude PI in the absence of detergent (Fig. S2C); GFP^+^ *TPA* staining for PI following antibody incubation (see Fig. S5 for gating strategies for individual antisera), therefore, would indicate organisms with disrupted OMs (Fig. S2B). As shown in Fig. 7B-C, in the absence of antibodies, <1% of *in vitro*-cultivated GFP^+^ *TPA* were PI^+^. We also saw negligible staining with PI when GFP^+^ *TPA* were co-cultivated with either NRS (1.14%) or α-TP0751 antibodies (4.57%). In contrast, co-cultivation with two independent immune sera obtained from rabbits infected with the Nichols strain (IRS Nic-1 and Nic-2) led to a marked increase in PI^+^ treponemes (∼45-52%; *p* ≤ 0.0001) compared to both NRS and α-TP0751 (Fig. 7B-C). The effect was even more pronounced (∼96% of treponemes were PI^+^) when GFP^+^ *TPA* were co-cultivated with α-BamA ECL4 antibodies (Fig. 7B-C). Interestingly, two independent IRS obtained from rabbits infected with the SS14 strain (IRS SS14-1 and SS14-2) also decreased the growth of GFP^+^ *TPA* Nichols, but to a lesser extent than the Nichols-specific IRS (approximately one-log_10_; Fig. 7A) and with a comparatively modest increase in PI labeling (∼12-18%) (Fig. 7B-C). Given that both IRS SS14-1 and SS14-2 strongly inhibited growth of the *TPA* SS14 strain (41), the decreased effectiveness of these sera against the heterologous strain likely reflects OMP variability between these two *TPA* reference strains (42–44). However, it also is possible that the SS14 IRS used in these studies are less effective at inhibiting growth compared to their Nichols counterparts.

**Fig 7.**
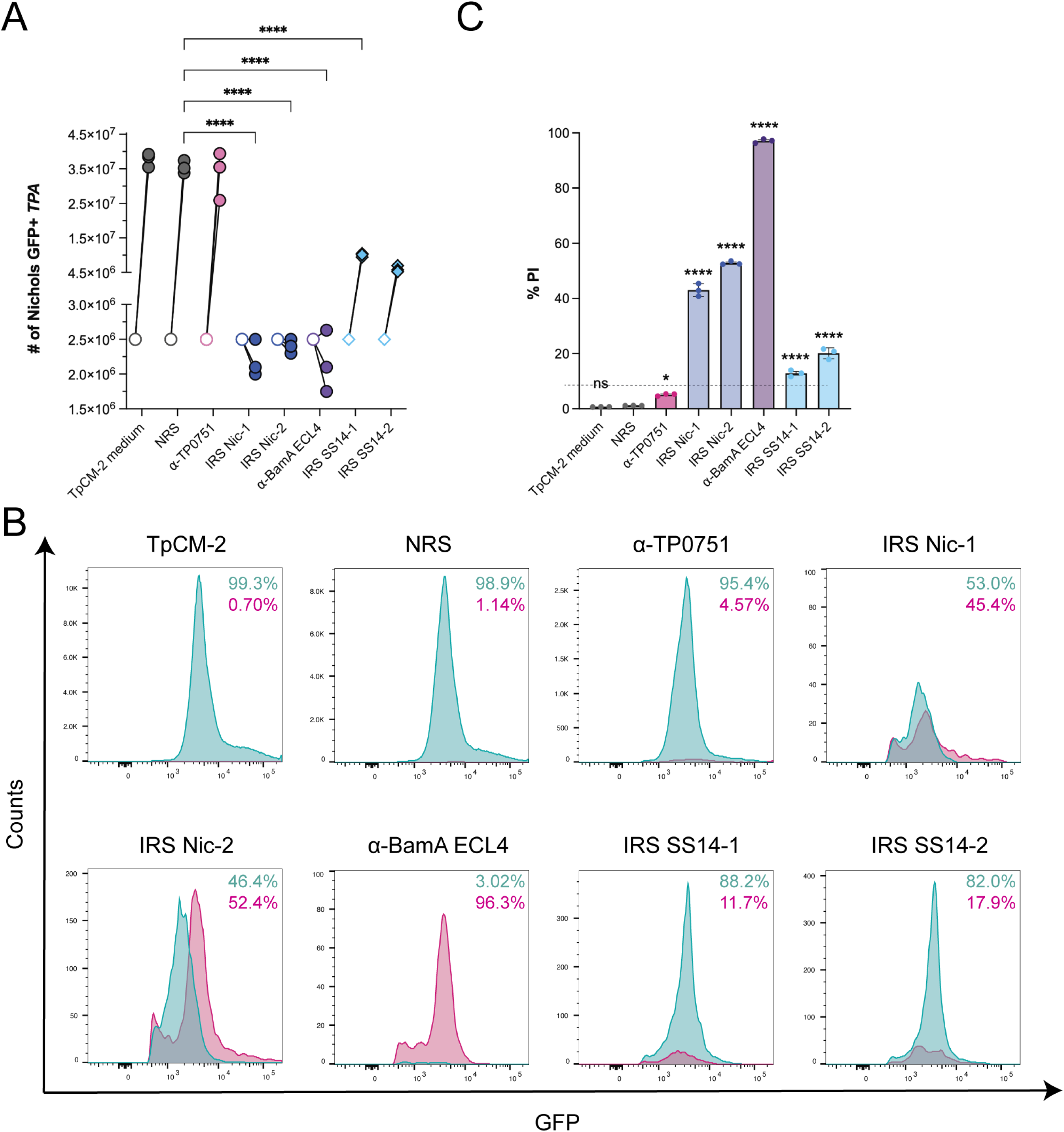
Disruption of *TPA* outer membranes by immune rabbit sera and rabbit anti-BamA ECL4. (**A**) Enumeration of GFP^+^ *TPA* co-cultured *in vitro* with Sf1Ep cells in the presence of normal rabbit serum (NRS), immune rabbit serum (IRS) from two Nichols immune rabbits (IRS Nic-1 and IRS Nic-2) or two SS14 immune rabbits (IRS SS14-1 and IRS SS14-2), and rabbit antisera against either TP0751 or BamA ECL4. Symbols indicate the spirochete densities before (open) and seven days after (closed) the addition of antisera. (**B**) Flow cytometry histograms portraying the percentages of PI^-^ (cyan) and PI^+^ (magenta) organisms gated from GFP^+^ populations incubated with the indicated sera at a final concentration of 10% (also see Fig. S5). (**C**) Percentage of PI^+^ organisms within each GFP^+^ population. Bars represent the mean ± standard deviation for three biological replicates per condition. *, *p* ≤ 0.05; or \*\*\*\**p* ≤ 0.0001 compared to NRS.

## Discussion

The refinement of systems for the long-term cultivation *in vitro* (12, 39) and genetic manipulation (13, 45) of *TPA* has ushered in a new era for syphilis research. Importantly, the ability to transform *TPA* enabled targeted mutagenesis (13, 14) as well as, most recently, the generation of a fluorescent *TPA* SS14 isolate (21). In this study, we generated a GFP^+^ *TPA* Nichols isolate that replicated at wild-type levels *in vitro*, was fully virulent in rabbits infected by intratesticular and intradermal inoculation, and disseminated to distal sites at levels comparable to the parental strain. We then used this strain to assess functional antibodies within syphilitic serum generated during experimental infection with *TPA* and following immunization with a protein scaffold containing *TPA* Nichols BamA ECL4, a known opsonic target (37, 41). Our findings illustrate the broad utility of fluorescent reporters for dissecting host-pathogen interactions during syphilitic infection, including antibody-mediated damage to the spirochetal OM.

Previous studies have shown that *TPA* adheres to a broad range of host cells, including epithelial, fibroblast, and endothelial cells (46–51). Preincubation of *TPA* with IRS and human syphilitic sera blocks spirochete attachment to host cells and/or extracellular matrix components, implying the existence of specific surface adhesins (46–52). Short- and long-term *in vitro* cultivation of *TPA*, an extreme auxotroph, also requires continuous intimate contact with host cells (11, 12), presumably to scavenge host-derived nutrients. An important question, therefore, is whether *TPA* is exclusively an extracellular pathogen within its obligate human host and whether intracellular residence is part of its strategy for persistence and immune evasion; indeed, sightings of ostensibly viable treponemes within Sf1Ep and other nonphagocytic cells have been reported over the years (25, 53, 54). Herein we showed the utility of GFP^+^ *TPA* for assessing cellular interactions by live spirochetes *in vitro* and *in vivo*. Consistent with *TPA* being predominantly an extracellular pathogen (55), confocal imaging of GFP^+^ *TPA* co-cultured with Sf1Ep cells and within testes from infected rabbits revealed numerous surface-attached spirochetes and no evidence of intracellular organisms. Vascular escape by *TPA* is a critical step in hematogenous dissemination and target organ invasion (2, 56). Classic electron microscopy studies showed *TPA* migrating through the intercellular junctions separating human umbilical vein endothelial cells, a process designated ‘inter-junctional penetration’ (48, 57), a finding also consistent with the spirochete’s extracellular lifestyle. The concept that OMPs can have both physiological and virulence-related functions is well established (58–61) and can be applied to *TPA* now that its repertoire of OMPs – the *TPA* OMPeome – has been delineated (34). Live imaging of GFP^+^ *TPA* should help elucidate how syphilis spirochetes penetrate tissues and, in concert with mutagenesis, help identify the responsible OMP culprits. Recently, we generated evidence that antibodies directed against specific ECLs of the spirochete’s FadL fatty acid transporters prevent their attachment to Sf1EP cells (41). Blockage of attachment of GFP^+^ *TPA* by antibodies against ECLs provides an additional means of assessing the contribution(s) of individual *TPA* OMPs to adherence and dissemination, information of great value for vaccine design as well as unraveling of key events during the disease process.

Antibody-mediated clearance by macrophages is thought to be crucial for controlling syphilitic infection (10, 35). While there currently is no ‘true’ correlate of protection for syphilis in humans, *ex vivo* opsonophagocytosis assays using sera from experimentally-infected animals and human syphilitic sera are regarded as a surrogate of protection (36, 37, 41, 62–64). Analysis of the molecular architecture of the *TPA* OM indicates that the presumptive targets for opsonic antibodies in immune sera reside largely within the OMPeome (9, 30, 34, 35, 65, 66). Previously, we demonstrated that antibodies against ECLs of several OMPs, including BamA ECL4, are strongly opsonic in assays using rabbit peritoneal macrophages and murine bone marrow-derived macrophages (37, 41). Using GFP^+^ *TPA,* we streamlined this assay by eliminating the need for antibody labeling to detect surface-bound versus internalized organisms. Understanding *TPA* interactions with macrophages is also directly relevant for efforts to deconvolute the spirochete’s strategies for ‘stealth pathogenicity’ (9). It has long been known that only a subset of organisms incubated with IRS or antibodies to individual OMPs are susceptible to internalization by macrophages, leading to the assumption that there is heterogeneous surface antigenicity within spirochete populations (63, 65, 67). The need for surface-directed antibodies to promote macrophage uptake theoretically provides a window during early infection during which *TPA* can disseminate and invade while bypassing innate immune pathogen surveillance systems (68). Furthermore, studies by our group have shown that, because of the lack of lipopolysaccharide and the paucity of lipoproteins on the spirochetal surface (35), macrophage activation requires internalization and degradation of *TPA* within phagocytic vacuoles, thereby liberating PAMPs for binding to Toll-like receptors lining the vacuole (36, 69). GFP^+^ *TPA* represents an important addition to the armamentarium for studying the underlying cell and immunobiology of this enigmatic disease.

Staining with the impermeant dye PI provided a straightforward means of using flow cytometry to assess OM integrity as a quantifiable presumptive marker for spirochete killing during incubation with IRS and anti-ECL antibodies. Flow cytometric analyses of GFP^+^ *TPA* co-cultivated in the presence of heat-inactivated IRS or BamA ECL4-specific antisera demonstrated that both appear to exert their bactericidal effects by damaging the spirochete’s fragile OM by a complement- and Fc-receptor-independent mechanism. Cryoelectron microscopy has shown that antibodies against ECL4 of *E. coli* BamA freeze the BAM complex in the ‘open’ conformation, preventing insertion of newly synthesized OMPs into the OM bilayer (70). It is reasonable to presume that interference with OM biogenesis is responsible for the loss of OM integrity and resultant potent bactericidal effect of *TPA* BamA ECL4 antibodies observed herein. BamA antibodies also may be a major contributor to the killing capacity of IRS. Importantly, the substantively greater capacity for OM disruption by anti-BamA ECL4 clearly argues for the importance of including this antigen in a syphilis vaccine cocktail. Studies in the 1970s using IRS to passively immunize rabbits against intradermal inoculation revealed that IRS is protective, but that *TPA* replication and lesion formation resumed once serum administration was discontinued (71, 72). It is tempting to speculate that the *in vitro* observations made herein reflect the suppressive effects of immune sera reported in these classic studies. In addition to providing a novel analytic tool to clarify *TPA*’s enigmatic interactions with surface-directed antibodies, the PI-GFP flow cytometric assay provides a means for delineating the protective mechanisms of an ECL antiserum as well as the ability of organisms to evade its growth inhibitory and/or killing effects, which could be clinically significant as antibody titers decline following vaccination.

While rabbits infected with *TPA* develop complete immunity to homologous challenge, protection against heterologous isolates is less robust (73). Consistent with these and other studies (41), we observed significantly less growth inhibition and OM damage when the GFP^+^ *TPA* Nichols strain was co-cultivated with heterologous IRS generated against the SS14 strain. We interpret these results to indicate the importance of *TPA* strain-specific antibodies for maximal bactericidal activity. Evidence for sequence variability within OMPs encoded by *TPA* clinical isolates is well documented (37, 41, 44, 74–78). Even single nonsynonymous amino acid substitutions in ECL4 of BamA can abrogate antibody binding (38). Substitutions in ECL3 from the FadL ortholog TP0865 also markedly affected antigenicity in SS14 compared to the Nichols (41). In a clinical scenario, antibodies targeting conserved ECLs likely provide a measure of cross-immunity, while OMP variability provides a means for immune evasion by organisms circulating within at-risk human populations. How these two factors balance out is a major issue requiring further investigation. Phenotypic characterization of isogenic GFP^+^ *TPA* strains expressing OMP ECL variants could help determine how sequence variability impacts clearance by functional antibodies in IRS and antisera from ECL-immunized animals, facilitating screening to identify ECL combinations capable of protecting against heterologous strains.

## Methods

### Ethics Statement

Animal experiments were conducted following the *Guide for the Care and Use of Laboratory Animals* (8th Edition) in accordance with protocols reviewed and approved by the UConn Health Institutional Animal Care and Use Committee (AP-201085) under the auspices of Public Health Service assurance number A3471-01 (D16-00295).

### Routine propagation of *TPA* in rabbits

*TPA* Nichols was propagated by intratesticular inoculation of adult male New Zealand White (NZW) rabbits as previously described (64, 65). Treponemes were harvested at peak orchitis in 0.5 - 1 ml of CMRL medium (ThermoFisher) supplemented with 10% heat-inactivated normal rabbit sera (NRS). After 2 hrs, treponemes were recovered and enumerated by darkfield microscopy (DFM) using a Petroff-Hausser counting chamber (Hausser Scientific, Horsham, PA, USA).

### Generation of immune rabbit and mouse syphilitic sera

Immune rabbit sera (IRS) against *TPA* Nichols and SS14 strains were generated by inoculation of rapid plasma reagin-nonreactive adult male NZW rabbits (n = 2 per strain) in each testis with 1 × 10^7^ treponemes in 0.5 ml of CMRL supplemented with 10% NRS. The immune status of each rabbit was confirmed 60 days post-inoculation by intradermal challenge with 1 × 10^3^ freshly extracted *TPA* of the same strain at each of eight sites on their shaved backs. Animals were euthanized and exsanguinated once their immune status had been confirmed by lack of lesion development. Animal identification numbers (ID#) for IRS Nic-1 and Nic-2 were ID#112 and ID#717, respectively. Animal IDs for IRS SS14-1 and SS14-2 were ID#759 and ID#761, respectively. To generate mouse syphilitic sera (MSS), five male and five female 6- to 8-week-old C3H/HeJ mice were inoculated intradermally, intraperitoneally, intrarectally, and intra-genitally with a total of 1 x 10^8^ total organisms per animal as previously described (37, 79). Mice were sacrificed on day 84 post-inoculation and exsanguinated to create a pool of MSS.

### *In vitro* cultivation of *TPA*

*TPA* were co-cultured with cottontail rabbit epithelial cells (Sf1Ep) in *TPA* culture medium 2 (TpCM-2) under microaerophilic (MA) conditions as previously described (12, 39). Briefly, Sf1Ep were seeded at 2 × 10^4^ cells per well in a 24-well culture plate and incubated overnight at 37°C. The following day, wells were washed with TpCM-2 followed by the addition of fresh TpCM-2 medium and 2.5 × 10^6^ *TPA* were added per well. After 7 days and weekly thereafter, treponemes were harvested by trypsinization, enumerated by DFM, and passaged on to fresh Sf1Ep cells. The same procedure was used for *in vitro* cultivation in 6-well plates with the exception that Sf1Ep cells were seeded at 5 × 10^4^ cells per well and 5 x 10^6^ *TPA* were added per well.

### Routine DNA manipulation and cloning

*Escherichia coli* Stellar cells (TaKaRa, Mountain View, CA) were used for routine cloning and isolation of plasmid DNA. *E. coli* ClearColi strain (Research Corporation Technologies, Tucson, AZ) was used for isolation of lipopolysaccharide-free plasmid DNA. *E. coli* cultures were maintained in Lysogeny broth (LB) or LB agar supplemented with the appropriate antibiotics (ampicillin, 100 μg/ml and/or kanamycin, 50 μg/ml). Plasmid DNA was purified from *E. coli* using QIAprep kits (Qiagen, Valencia, CA, USA) according to manufacturer’s instructions. Synthetic gene fragments were purchased from Integrated DNA Technologies, Inc. (Coralville, IA, USA). *TPA* genomic DNA was extracted using the DNeasy Blood and Tissue kit (Qiagen). Oligonucleotide primers (Table S1) were purchased from Sigma-Aldrich (St. Louis, MO, USA). Routine cloning was performed using the In-Fusion HD Cloning Plus kit (Takara Bio USA, Inc., Mountain View, CA, USA). Routine and high-fidelity PCR amplifications were performed using RedTaq (Denville Scientific, Metuchen, NJ, USA) and CloneAmp HiFi (Takara Bio USA, Inc.), respectively. Plasmid constructs were confirmed by Sanger sequencing (Azenta Life Sciences, South Plainfield, NJ, USA) using primers listed in Table S1 and analyzed using MacVector (MacVector, Inc., Cary, NC, USA).

### Construction of a suicide vector for replacement of *tprA* with a GFP cassette

A suicide vector used for chromosomal insertion of a cassette containing extra-superfolder GFP and a kanamycin-resistance marker (*kanR*) was generated by cloning a ∼4.3 kb amplification product containing *tprA* (*tp0009*) plus flanking DNA into BamHI-digested pUC19 using primers tprA-FW and tprA-RV (Table S1 and Fig. S1A). The *tprA* coding sequence was then replaced with a fragment containing codon-optimized *kanR* from *Proteus vulgaris* (80) under the control of the *tpp47* (*tp0574*) promoter and ribosomal binding site (RBS) (pMC5722; Fig. S1A). A synthetic fragment containing a codon-optimized version of extra-superfolder GFP (22, 24) under the control of the *flaA1* (*tp0249*) promoter and RBS (81) was then inserted upstream of *kanR* (pMC5836; Fig. S1B).

### Generation of fluorescent *TPA*

A *TPA* Nichols strain constitutively expressing GFP was generated by transforming *in vitro*-cultivated treponemes with 15 μg of lipopolysaccharide-free pMC5836 plasmid DNA (Fig. S1A) as previously described (13, 45). Briefly, *TPA* co-cultured with Sf1Ep cells for 1-week were trypsinized to release bound treponemes. ∼5 × 10^7^ organisms were transferred to one well of a 24-well plate seeded the prior day with 2 × 10^4^ Sf1Ep cells. The total volume in each well was brought to 2.5 ml with pre-equilibrated TpCM-2 and returned to the incubator. After 2 days, 1 ml of spent media was removed and replaced with 1 ml of fresh pre-equilibrated TpCM-2. On day 4, spent media was removed gently and replaced with 0.5 ml of Transformation Buffer (50 mM CaCl_2_, 10 mM Tris pH 7.4) containing 15 μg of LPS-free plasmid DNA. Following a 10 min incubation at 34°C under MA conditions, cells were washed twice gently with pre-equilibrated TpCM-2 and recovered overnight in 2.5 ml of fresh TpCM-2 at 37°C under MA conditions. The following day, kanamycin (200 μg/ml final concentration; Sigma-Aldrich, St. Louis, MO, USA) was added and plates were returned to the incubator for 48 hours. Following incubation, the media was exchanged with fresh TpCM-2 containing 200 μg/ml kanamycin (TpCM-2Kan) per well. After two weeks, *TPA* were trypsinized, enumerated by DFM, and passaged to a new 24-well plate seeded with fresh Sf1Ep cells in TpCM-2Kan. Untreated, Transformation buffer (TB) alone, and “no kanamycin” control transformations were performed in parallel. Wells were examined biweekly by DFM and/or epifluorescence microscopy for motile treponemes. Once co-cultures reached a density of ∼3 × 10^7^ *TPA* per ml, cells were trypsinized, enumerated by DMF, and then passed into 6-well plates seeded with 5 × 10^4^ Sf1Ep cells per well. Allelic replacement of *tprA* with either *gfp*-*kanR* or *kanR* was confirmed by PCR amplification of genomic DNA using primers described in Table S1 and Fig. S1C-D. The resulting nonfluorescent (*kanR*) and GFP^+^ (*gfp-kanR)* strains were designated SRL001 and SRL002, respectively.

### Whole-genome sequencing

Genomic DNA was extracted from *in vitro*-cultivated GFP^+^ *TPA* collected at passages 6 (week 12), 9 (week 20) and 12 (week 26) as described above. For GFP^+^ *TPA* libraries were generated using Kapa Hyper Prep kit (Kapa Biosystems, Inc., Wilmington, MA, USA) and sequenced on an Illumina iSeq 100 system (Illumina, San Diego, CA, USA), utilizing 150-base paired-end reads. To determine if genetic engineering induced novel mutations, we also sequenced the WT parent used to generate the GFP^+^ strain, designated ‘Nichols-Farmington’, using DNA extracted from infected testes tissue, mixed with purified human DNA at 1:99 ratio of total DNA as previously described (82). Libraries were prepared using SureSelect XT HS target enrichment system (Agilent Technologies, Santa Clara, CA) with sequencing performed on the Illumina MiSeq system (Illumina, San Diego, CA, USA) as previously described (83). Raw read data were analyzed using a modified version of the bioinformatic pipeline available at https://github.com/IDEELResearch/tpallidum_genomics. Quality control of the raw reads was performed using FastQC (84), followed by adapter trimming with Cutadapt (v4.4)(85). Reads with a sequence quality score ³20 were retained. Processed reads were aligned to the NCBI Nichols strain reference genome (CP004010.2) (42, 86) using minimap2 (v2.26) (87). Reads mapping to the highly variable *tprK* locus (14, 88–90) were eliminated from our analyses. Post-alignment filtering and variant calling were conducted with SAMtools and BCFtools (v1.9) (91). Transgenic insert breakpoints were identified using Gridds (v2.13.2) (92). Statistical analysis and visualization were carried out using R (v4.3.3) (93) and ggplot2 (v3.5.1) (94). Raw read data for all passages of the GFP^+^ and WT Nichols-Farmington *TPA* were deposited in the Sequence Read Archive database (BioProject PRJNA1134712).

### Flow cytometric comparison of GFP expression by GFP^+^ *TPA* cultivated *in vitro* and recovered from rabbits

∼2 × 10^8^ *in vitro*-cultivated GFP^+^ *TPA* were harvested from a 6-well plate as described above. For comparison with *in vivo*, ∼4 × 10^8^ GFP^+^ *TPA* were harvested from rabbit testes at peak orchitis as described above. Non-fluorescent WT *TPA* Nichols grown under the same conditions were included as a negative control. Treponemes were pelleted at 8,000 × *g* for 10 minutes, washed twice with phosphate-buffered saline (PBS), and fixed for 10 min at 4°C in PBS containing 2% paraformaldehyde and 0.001% PI (1.5 μM final conc.) in the presence or absence of 0.01% Triton X-100. After fixation, treponemes were pelleted at 8,000 × *g* for 10 minutes, washed twice with PBS, and the resulting pellet was resuspended in 200 µl of PBS and transferred to a 96-well plate; 25 μl of each sample was analyzed on a FACSymphony A5 SE flow cytometer (BD Biosciences, Franklin Lakes, NJ, USA). Data were analyzed using FlowJo Version 10.7.1 software (BD Biosciences). The mean fluorescence intensity (MFI) and percentages (%) of PI^+^ and/or GFP^+^ events were calculated from data filtered to exclude non-spirochetal (PI/GFP double negative) events. Details regarding the gating strategy used for analysis of flow cytometry data are presented in Fig. S2.

### Virulence testing of GFP^+^ *TPA* by intratesticular inoculation

Infectivity of *in vitro*-cultivated WT and GFP^+^ *TPA* was compared by intratesticular inoculation of NZW rabbits as described above (see Routine propagation of *TPA* in rabbits). At peak orchitis, animals were sacrificed and spirochete viability and burdens in testes determined by DFM. WT and GFP^+^ *TPA* harvested from the first rabbit (Rabbit 1) were used to inoculate a second NZW rabbit (Rabbit 2). At peak orchitis, the second rabbit for each strain was sacrificed and spirochete viability and burdens in testes determined by DFM. Popliteal lymph nodes (LNs; two per rabbit, per strain, per experiment) and whole blood (three 1-ml aliquots per rabbit, per strain, per experiment) were collected from Rabbit 2 at the time of sacrifice to assess dissemination by qPCR. Individual LNs were placed in 200 μl of DNA/RNA Shield (Zymo), while each 1 ml blood sample was mixed with 2 mls of DNA/RNA Shield (Zymo). Samples were stored at -20°C until extraction using the DNeasy Blood and Tissue kit (Qiagen). qPCR assays for *polA* and rabbit β-actin were performed as previously described (40, 63). WT and GFP^+^ *TPA* were compared in three independent serial passage experiments. Statistical analyses were conducted using Prism (v. 9.5.1; GraphPad Software, San Diego, CA, USA). An unpaired two-tailed *t*-test was used to compare the number of WT and GFP^+^ *TPA* burdens. A *p*-value of ≤ 0.05 was considered statistically significant.

### Assessment of infectivity of GFP^+^ *TPA* by intradermal challenge

Serial dilutions of *in vitro*-cultivated WT and GFP^+^ *TPA* (10^5^ – 10^1^ treponemes per site) in CMRL with 10% NRS were used to inoculate each side of the shaved backs of male NZW rabbits. Animals were examined daily to monitor the development, morphologic appearance, and progression of lesions. Lesions were measured daily with digital calipers beginning 7 days p.i. until sacrifice on day 30. After euthanasia, the dorsal skin was depilated to remove any remaining fur, cleaned with 70% ethanol, and cutaneous lesions were excised using a 4 mm punch biopsy tool for cryosectioning (described below) and qPCR. Tissues were placed in DNA/RNA Shield (Zymo) and stored at -20°C until extraction using the DNeasy Blood and Tissue kit (Qiagen). qPCR assays for *polA* and rabbit β-actin were performed as previously described (40, 63). Statistical analyses were conducted using Prism (v. 9.5.1; GraphPad Software, San Diego, CA, USA). A repeated measures two-tailed ANOVA was used to compare lesion circumferences at each time point for rabbits inoculated with the same dose of either WT or GFP^+^ *TPA.* A paired *t*-test was used to compare spirochete burdens for WT and GFP^+^ *TPA* in lesions for 10^5^ and 10^4^ inocula. Bonferroni’s correction for multiple comparisons was applied and *p*-values ≤ 0.05 were considered significant.

### Confocal imaging of GFP^+^ *TPA* co-cultured with Sf1Ep rabbit epithelial cells

Round glass cover slips (13 mm) were autoclaved and transferred to a six-well culture plate seeded with 5 × 10^4^ Sf1Ep cells per well and then incubated overnight at 37°C. The following day, wells were washed with TpCM-2 and 5 × 10^6^ GFP^+^ *TPA* in TpCM-2 added per well. After 7 days at 37°C under MA conditions, cells were washed twice with PBS then fixed for 15 min with 4% paraformaldehyde in PBS. After fixation, cells were washed with TpCM-2 as described above and cover slips transferred to clean 6-well plate containing 2 ml of 1X PBS per well. Sf1Ep cell membranes were stained with Cholera Toxin AF647 (200 ng/ml final conc; ThermoFisher) (95) for 30 min followed by staining of host cell nuclei with 4′,6-diamidino-2-phenylindole (DAPI, 5 μg/ml final conc.; ThermoFisher) for 10 min. After two washes with PBS, coverslips were transferred cell side down onto a clean slide containing VECTASHIELD mounting medium (Vector Laboratories, Inc.) and sealed with nail polish. Individual 1 μm optical sections were acquired using a Zeiss 880 confocal microscope equipped with 63X/1.4 Plan-Apochromat oil objective and processed using ZEN3.5 Blue (Carl Zeiss Microscopy, White Plains, NY, USA).

### Confocal imaging of GFP^+^ *TPA* in infected tissues

GFP^+^ *TPA*-infected testis tissue (∼0.5 – 1 cm) was fixed in 2% paraformaldehyde for 1 hr at 4°C and then washed at least three times with PBS for 10 min at room temperature. For cryosectioning, fixed testis tissues were transferred to 15% sucrose in PBS for 6-12 hrs, followed by overnight incubation at 4°C in 20% sucrose in PBS before being embedded in OCT compound using a 2-methyl-butane/dry ice/ethanol bath. Embedded tissues were stored at -80°C until sectioning. 7 μm sections were cut using a Leica CM3050 S cryostat. Prior to imaging, sections were incubated for 30 min with Cholera Toxin AF647 (200 ng/ml final conc.; ThermoFisher) (95), washed briefly with PBS containing 0.05% Tween 20 (PBST) followed by incubation with DAPI (5 μg/ml final conc.; ThermoFisher) for 10 min. Slides were washed thoroughly three times with PBST, rinsed with deionized water, and allowed to air dry. Sections were preserved in VECTASHIELD mounting medium (Vector Laboratories, Inc.), sealed with a coverslip. Individual 1 μm optical sections were acquired using a Zeiss 880 confocal microscope equipped with a with 63x/1.4 Plan-Apochromat oil objective. Images were processed using ZEN3.5 Blue.

### Opsonophagocytosis assay of GFP^+^ *TPA* by murine bone marrow-derived macrophages

Bone marrow-derived macrophages were generated from C3H/HeJ mice as previously described (37, 41), plated at a final concentration of 1 × 10^5^ cells per well in Millicell EZ 8-well chamber slides (Sigma-Aldrich), and incubated overnight at 37°C. The following day, the medium was replaced with fresh Dulbecco’s Modified Eagle Medium (DMEM) supplemented with 10% FBS prior to the addition of *TPA*. For opsonophagocytosis assays, *in vitro-*cultivated GFP^+^ treponemes were harvested using Dissociation media as previous described (39). For opsonization, ∼1 × 10^6^ *TPA* in 250 μl was preincubated at RT for 2 hr in DMEM supplemented with 1:10 dilutions of mouse syphilitic sera (MSS) generated against *TPA* Nichols (37) or mouse antisera directed against *TPA* Nichols BamA ECL4 (38). Following preincubation, all conditions with or without sera were added to cells for 4 h at 37°C and an MOI of 10:1. Negative controls included normal mouse sera (NMS) and mouse antisera directed against TP0751 (40). Following a 4 h incubation, supernatants were removed and macrophages were fixed/permeabilized with 2% paraformaldehyde and 0.01% Triton X-100 for 10 min at RT. Each well was rinsed with PBS and blocked with 1% bovine serum albumin in PBS overnight at 4°C. Wells were then incubated with Cholera Toxin AF647 (200 ng/ml final conc; ThermoFisher) for 30 min and DAPI (5 μg/ml final conc; ThermoFisher) for 10 min and washed thoroughly three times with PBST, rinsed with deionized water to remove salt, and allowed to air dry. Samples were preserved in VECTASHIELD mounting medium (Vector Laboratories, Inc., Newark, CA, USA), sealed with a coverslip. Internalization of *TPA* was assessed in a blinded fashion by acquiring epifluorescence images of at least 100 macrophages per well using an Olympus BX-41 microscope equipped with a 40x/1.00 UPlan-Apochromat oil iris objective. Images were processed with VisiView (v. 5.0.0.7; Visitron Systems GmbH). Assays were performed in triplicate for each condition tested. The phagocytic index for each sample was calculated by dividing the number of internalized spirochetes by the total number of cells imaged and multiplying by 100. Confocal images (12-15 1-μm optical sections) were acquired using a Zeiss 880 equipped with a 63x/1.4 Plan-Apochromat oil objective and processed using ZEN3.5 Blue. Statistical analyses were conducted using Prism (v. 9.5.1; GraphPad Software, San Diego, CA, USA). One-way ANOVA was used to compare phagocytic indices using Newman-Keuls and Bonferroni’s correction for multiple comparisons, respectively. *p*-values ≤ 0.05 were considered significant.

### Flow cytometric assessment of growth inhibition and OM disruption during incubation with IRS and antibodies to BamA ECL4

Sf1Ep cells were cultured in a 24-well plate ON as described above. 2.5 × 10^6^ freshly harvested GFP^+^ *TPA* were added to each well along with 10% NRS, IRS from two Nichols immune rabbits (IRS Nic-1 and IRS Nic-2), or IRS from two SS14 immune rabbits (IRS SS14-1 and IRS SS14-2), or rabbit polyclonal antisera against *TPA* Nichols BamA ECL4 (37, 38) or TP0751 (40) and then incubated under MA conditions. All assays were performed in triplicate. After seven days, spent media was transferred to a clean centrifuge tube, cells were washed once with 200 μl of trypsin-EDTA (Sigma-Aldrich), and the recovered material was combined with the spent media from the same well. An additional 200 μl of trypsin-EDTA was then added to each well and incubated at 34°C for 5 min to release treponemes. Following trypsinization, the combined material from each well was centrifuged at 130 × *g* for 5 min and the number of total treponemes recovered enumerated by DFM as described above. The combined material from each well was processed for flow cytometry and analyzed as described above. The percentage of PI^+^ organisms within the GFP^+^ population was determined after excluding non-spirochetal (*i.e*., double negative) events as shown in Fig. S2. Statistical analyses were conducted using Prism (v. 9.5.1; GraphPad Software, San Diego, CA, USA). A two-way ANOVA and a one-way ANOVA were used to compare *TPA* growth *in vitro*, and % PI staining with Tukey correction for multiple comparisons, respectively. *p*-values ≤ 0.05 were considered significant.

## Acknowledgments

We thank Morgan LeDoyt and Kemar Edwards for outstanding technical support. The authors would like to acknowledge support from Dr. Evan Jellison (UConn Health Flow Cytometry Core Facility), Dr. Zhifang Hao (UConn Health Research Histology Core), Susan Staurovsky (UConn Health Richard D. Berlin Center for Cell Analysis and Modeling) and Rachel Spreng (Duke Human Vaccine Institute).

## Author contributions

Conceptualization: K.N.D., J.D.R, K.L.H and M.J.C.

Methodology: K.N.D., C.F.V., C.M.H., K.P.C., F.A., J.B.P., K.L.H and M.J.C.

Investigation, K.N.D., C.F.V., C.M.H, F.A., K.L.H and M.J.C.

Writing – Original Draft, K.N.D., J.D.R, K.L.H and M.J.C; all authors reviewed and edited the manuscript.

Supervision: K.L.H and M.J.C.

Funding Acquisition, K.N.D., J.D.R., K.L.H, and M.J.C.

## Supplemental Material

**Fig S1. Replacement of *tprA* with extra-superfolder GFP and/or a kanamycin resistance gene.** Plasmids used to replace *tprA* in the *TPA* Nichols chromosome with a constitutively expressed codon-optimized extra-superfolder green fluorescent protein (*gfp*) transgene under the control of the *flaA1* promoter and a kanamycin-resistance gene (*kanR*) from *Proteus mirabilis* under the control of the *tpp47* promoter (pMC5836) (**A**) or the *kanR* cassette alone (**B**). Schematics showing the chromosomal regions used to insert the *gfp-kanR* (**C**) and *kanR* (**D**) cassettes. Arrows are used to indicate locations of primers (Table S1) used to confirm insertion. The expected size in base pairs (bp) for each amplicon is indicated above the corresponding line. Agarose gel images showing the corresponding PCR amplicons obtained using genomic DNA from GFP^+^ (**C**), *kanR* (**D**) *TPA* Nichols strains are shown below. Genomic DNA from WT *TPA* was used as a negative (no insert) control. Neg, No DNA control. DNA ladder (bp) is shown on the left of gels in **C** and **D**.

**Fig S2. Gating strategies used for flow cytometric analysis of WT and GFP^+^ *TPA*.** (**A**) Flow cytometry panels for unstained WT *TPA* used to define and exclude non-spirochetal (*i.e.*, double-negative) events. (**B**) Flow cytometry panels for detergent-treated (+ Triton), *in vitro*-cultivated WT *TPA* stained with PI used to define the spirochete population. (**C**) Flow cytometry panels for, *in vitro*-cultivated GFP^+^ *TPA* stained with PI in the absence of detergent (-Triton) used to define the GFP^+^ population and confirm exclusion of PI by intact treponemes. Results are representative of three independent experiments.

**Fig S3. Whole-genome sequencing to confirm replacement of *tprA* with *gfp*-*kanR* in GFP^+^ *TPA***. Genome sequencing confirms the replacement of *tprA* with the *gfp-kanR* cassette in GFP^+^ *TPA*. Assembled reads for *tprA* and flanking regions from GFP^+^ *TPA* after 6, 9 and 12 passages *in vitro* (**A-C**) and serial passages in rabbit testes (**D**, **E**) mapped against the *TPA* Nichols reference genome (left panels) and modified genome containing the *gfp-kanR* transgenes in place of *trpA* (right panels). The gaps in coverage in A-E left panels demonstrate complete replacement of the native *trpA* coding sequence in GFP^+^ *TPA*.

**Fig S4. Rabbit intradermal inoculations with GFP^+^ *TPA* mirrors lesion development of WT *TPA*.** Lesion circumferences measured in mm and averaged from rabbits (n = 3) inoculated intradermally with graded doses (1x10^4^ – 1x10^1^) of GFP^+^ and WT *TPA.* Lesions were measured beginning day 7 p.i. until sacrifice (day 30 p.i.). Values represent the mean ± standard deviation for three biological replicates per condition. **, *p* ≤ 0.01; or \*\*\**p* ≤ 0.001.

**Fig S5. Gating strategy used to assess OM disruption of *in vitro*-cultivated GFP^+^ *TPA*.** Flow cytometric panels used to exclude non-spirochetal (*i.e*., double negative) events and then assessing the percentage of PI^+^ organisms within the GFP^+^ population for each serum. Results are representative of three independent experiments.

**Video S1. Epifluorescence video showing motility of *in vitro*-cultivated GFP^+^ *TPA* Nichols strain.**

**Video S2. Epifluorescence video showing motility of *kanR TPA* Nichols strain.**

**Video S3. Z-stack of individual 1 μm optical sections showing surface localization of *in vitro*-cultivated GFP^+^ *TPA* Nichols strain co-cultured with Sf1Ep rabbit epithelial cells.**

**Video S4. Epifluorescence video showing motility of GFP^+^ *TPA* Nichols strain harvested from rabbit testes.**

**Video S5. Epifluorescence video showing motility of WT *TPA* Nichols strain harvested from rabbit testes.**

**Video S6. Z-stack of individual 1 μm optical sections showing surface localization of GFP^+^ *TPA* Nichols strain harvested from rabbit testes.**

## REFERENCES

1. Peeling RW, Mabey D, Chen XS, Garcia PJ. 2023. Syphilis. Lancet 402:336–346.

2. Radolf JD, Tramont EC, Salazar JC. 2019. Syphilis (*Treponema pallidum*). *In* Mandell GL, Dolin R, Blaser MJ (ed), Mandell, Douglas and Bennett’s Principles and Practice of Infectious Diseases. Churchill Livingtone Elsevier.

3. Cooper JM, Sanchez PJ. 2018. Congenital syphilis. Semin Perinatol 42:176–184.

4. Wozniak PS, Cantey JB, Zeray F, Leos NK, Michelow IC, Sheffield JS, Wendel GD, Sanchez PJ. 2023. The mortality of congenital syphilis. J Pediatr 263:113650.

5. Kojima N, Klausner JD. 2018. An update on the global epidemiology of syphilis. Curr Epidemiol Rep 5:24–38.

6. Moseley P, Bamford A, Eisen S, Lyall H, Kingston M, Thorne C, Pinera C, Rabie H, Prendergast AJ, Kadambari S. 2024. Resurgence of congenital syphilis: new strategies against an old foe. Lancet Infect Dis 24:e24–e35.

7. CDC. 2024. National Overview of STIs, 2022. Accessed

8. McDonald R, O’Callaghan K, Torrone E, Barbee L, Grey J, Jackson D, Woodworth K, Olsen E, Ludovic J, Mayes N, Chen S, Wingard R, Johnson Jones M, Drame F, Bachmann L, Romaguera R, Mena L. 2023. Vital Signs: Missed opportunities for preventing congenital syphilis - United States, 2022. MMWR Morb Mortal Wkly Rep 72:1269–1274.

9. Radolf JD, Deka RK, Anand A, Smajs D, Norgard MV, Yang XF. 2016. *Treponema pallidum*, the syphilis spirochete: making a living as a stealth pathogen. Nat Rev Microbiol 14:744–759.

10. Lafond RE, Lukehart SA. 2006. Biological basis for syphilis. Clin Microbiol Rev 19:29–49.

11. Fieldsteel AH, Cox DL, Moeckli RA. 1981. Cultivation of virulent *Treponema pallidum* in tissue culture. Infect Immun 32:908–15.

12. Edmondson DG, Hu B, Norris SJ. 2018. Long-term *in vitro* culture of the syphilis spirochete *Treponema pallidum* subsp. *pallidum*. mBio 9:e01153–18.

13. Romeis E, Tantalo L, Lieberman N, Phung Q, Greninger A, Giacani L. 2021. Genetic engineering of *Treponema pallidum* subsp. *pallidum*, the syphilis spirochete. PLoS Pathog 17:e1009612.

14. Romeis E, Lieberman NAP, Molini B, Tantalo LC, Chung B, Phung Q, Avendano C, Vorobieva A, Greninger AL, Giacani L. 2023. *Treponema pallidum* subsp. *pallidum* with an artificially impaired TprK antigenic variation system is attenuated in the rabbit model of syphilis. PLoS Pathog 19:e1011259.

15. Melican K, Richter-Dahlfors A. 2009. Real-time live imaging to study bacterial infections *in vivo*. Curr Opin Microbiol 12:31–6.

16. Konjufca V, Miller MJ. 2009. Two-photon microscopy of host-pathogen interactions: acquiring a dynamic picture of infection *in vivo*. Cell Microbiol 11:551–9.

17. Chaconas G, Moriarty TJ, Skare J, Hyde JA. 2021. Live Imaging. Curr Issues Mol Biol 42:385–408.

18. Bockenstedt LK, Gonzalez DG, Haberman AM, Belperron AA. 2012. Spirochete antigens persist near cartilage after murine Lyme borreliosis therapy. J Clin Invest 122:2652–60.

19. Dunham-Ems SM, Caimano MJ, Pal U, Wolgemuth CW, Eggers CH, Balic A, Radolf JD. 2009. Live imaging reveals a biphasic mode of dissemination of *Borrelia burgdorferi* within ticks. J Clin Invest 119:3652–65.

20. Caimano MJ, Groshong AM, Belperron A, Mao J, Hawley KL, Luthra A, Graham DE, Earnhart CG, Marconi RT, Bockenstedt LK, Blevins JS, Radolf JD. 2019. The RpoS gatekeeper in *Borrelia burgdorferi*: An invariant regulatory scheme that promotes spirochete persistence in reservoir hosts and niche diversity. Front Microbiol 10:1923.

21. Grillova L, Romeis E, Lieberman NAP, Tantalo LC, Xu LH, Molini B, Trejos AT, Lacey G, Goulding D, Thomson NR, Greninger AL, Giacani L. 2024. Bright new resources for syphilis research: Genetically encoded fluorescent tags for *Treponema pallidum* and Sf1Ep cells. Mol Microbiol.

22. Choi JY, Jang TH, Park HH. 2017. The mechanism of folding robustness revealed by the crystal structure of extra-superfolder GFP. FEBS Lett 591:442–447.

23. Pedelacq JD, Cabantous S, Tran T, Terwilliger TC, Waldo GS. 2006. Engineering and characterization of a superfolder green fluorescent protein. Nat Biotechnol 24:79–88.

24. Nagasundarapandian S, Merkel L, Budisa N, Govindan R, Ayyadurai N, Sriram S, Yun H, Lee SG. 2010. Engineering protein sequence composition for folding robustness renders efficient noncanonical amino acid incorporations. Chembiochem 11:2521–4.

25. Konishi H, Yoshii Z, Cox DL. 1986. Electron microscopy of *Treponema pallidum* (Nichols) cultivated in tissue cultures of Sf1Ep cells. Infect Immun 53:32–7.

26. Dunham-Ems SM, Caimano MJ, Eggers CH, Radolf JD. 2012. *Borrelia burgdorferi* requires the alternative sigma factor RpoS for dissemination within the vector during tick- to-mammal transmission. PLoS Pathog 8:e1002532.

27. Babb K, McAlister JD, Miller JC, Stevenson B. 2004. Molecular characterization of *Borrelia burgdorferi erp* promoter/operator elements. J Bacteriol 186:2745–56.

28. Bykowski T, Babb K, von Lackum K, Riley SP, Norris SJ, Stevenson B. 2006. Transcriptional regulation of the *Borrelia burgdorferi* antigenically variable VlsE surface protein. J Bacteriol 188:4879–89.

29. Miller JC, von Lackum K, Woodman ME, Stevenson B. 2006. Detection of *Borrelia burgdorferi* gene expression during mammalian infection using transcriptional fusions that produce green fluorescent protein. Microb Pathog 41:43–7.

30. Liu J, Howell JK, Bradley SD, Zheng Y, Zhou ZH, Norris SJ. 2010. Cellular architecture of *Treponema pallidum*: novel flagellum, periplasmic cone, and cell envelope as revealed by cryo electron tomography. J Mol Biol 403:546–61.

31. Izard J, Renken C, Hsieh CE, Desrosiers DC, Dunham-Ems S, La Vake C, Gebhardt LL, Limberger RJ, Cox DL, Marko M, Radolf JD. 2009. Cryo-electron tomography elucidates the molecular architecture of *Treponema pallidum*, the syphilis spirochete. J Bacteriol 191:7566–80.

32. Cox DL, Akins DR, Porcella SF, Norgard MV, Radolf JD. 1995. *Treponema pallidum* in gel microdroplets: a novel strategy for investigation of treponemal molecular architecture. Mol Microbiol 15:1151–64.

33. Edmondson DG, De Lay BD, Hanson BM, Kowis LE, Norris SJ. 2023. Clonal isolates of *Treponema pallidum* subsp. *pallidum* Nichols provide evidence for the occurrence of microevolution during experimental rabbit infection and *in vitro* culture. PLoS One 18:e0281187.

34. Hawley KL, Montezuma-Rusca JM, Delgado KN, Singh N, Uversky VN, Caimano MJ, Radolf JD, Luthra A. 2021. Structural modeling of the *Treponema pallidum* outer membrane protein repertoire: A road map for deconvolution of syphilis pathogenesis and development of a syphilis vaccine. J Bacteriol 203:e0008221.

35. Radolf JD, Kumar S. 2018. The *Treponema pallidum* outer membrane. Curr Top Microbiol Immunol 415:1–38.

36. Lukehart SA. 2008. Scientific monogamy: thirty years dancing with the same bug: 2007 Thomas Parran Award Lecture. Sex Transm Dis 35:2–7.

37. Ferguson MR, Delgado KN, McBride S, Orbe IC, La Vake CJ, Caimano MJ, Mendez Q, Moraes TF, Schryvers AB, Moody MA, Radolf JD, Weiner MP, Hawley KL. 2023. Use of Epivolve phage display to generate a monoclonal antibody with opsonic activity directed against a subdominant epitope on extracellular loop 4 of *Treponema pallidum* BamA (TP0326). Front Immunol 14:1222267.

38. Luthra A, Anand A, Hawley KL, LeDoyt M, La Vake CJ, Caimano MJ, Cruz AR, Salazar JC, Radolf JD. 2015. A homology model reveals novel structural features and an immunodominant surface loop/opsonic target in the *Treponema pallidum* BamA ortholog TP_0326. J Bacteriol 197:1906–20.

39. Edmondson DG, Norris SJ. 2021. *In vitro* cultivation of the syphilis spirochete *Treponema pallidum*. Curr Protoc 1:e44.

40. Luthra A, Montezuma-Rusca JM, La Vake CJ, LeDoyt M, Delgado KN, Davenport TC, Fiel-Gan M, Caimano MJ, Radolf JD, Hawley KL. 2020. Evidence that immunization with TP0751, a bipartite *Treponema pallidum* lipoprotein with an intrinsically disordered region and lipocalin fold, fails to protect in the rabbit model of experimental syphilis. PLoS Pathog 16:e1008871.

41. Delgado KN, Caimano MJ, Orbe IC, Vicente CF, La Vake CJ, Grassmann AA, Moody MA, Radolf JD, Hawley KL. 2024. Immunodominant extracellular loops of *Treponema pallidum* FadL outer membrane proteins elicit antibodies with opsonic and growth- inhibitory activities. Biorxiv 2024.07.30.605823.

42. Petrosova H, Pospisilova P, Strouhal M, Cejkova D, Zobanikova M, Mikalova L, Sodergren E, Weinstock GM, Smajs D. 2013. Resequencing of *Treponema pallidum* ssp. *pallidum* strains Nichols and SS14: correction of sequencing errors resulted in increased separation of syphilis treponeme subclusters. PLoS One 8:e74319.

43. Nechvatal L, Petrosova H, Grillova L, Pospisilova P, Mikalova L, Strnadel R, Kuklova I, Kojanova M, Kreidlova M, Vanousova D, Prochazka P, Zakoucka H, Krchnakova A, Smajs D. 2014. Syphilis-causing strains belong to separate SS14-like or Nichols-like groups as defined by multilocus analysis of 19 *Treponema pallidum* strains. Int J Med Microbiol 304:645–53.

44. Kumar S, Caimano MJ, Anand A, Dey A, Hawley KL, LeDoyt ME, La Vake CJ, Cruz AR, Ramirez LG, Pastekova L, Bezsonova I, Smajs D, Salazar JC, Radolf JD. 2018. Sequence variation of rare outer membrane protein beta-barrel domains in clinical strains provides insights into the evolution of *Treponema pallidum* subsp. *pallidum*, the syphilis spirochete. mBio 9:e01006–18.

45. Phan A, Romeis E, Tantalo L, Giacani L. 2022. *In vitro* transformation and selection of *Treponema pallidum* subsp. *pallidum*. Curr Protoc 2:e507.

46. Hayes NS, Muse KE, Collier AM, Baseman JB. 1977. Parasitism by virulent *Treponema pallidum* of host cell surfaces. Infect Immun 17:174–86.

47. Lee JH, Choi HJ, Jung J, Lee MG, Lee JB, Lee KH. 2003. Receptors for *Treponema pallidum* attachment to the surface and matrix proteins of cultured human dermal microvascular endothelial cells. Yonsei Med J 44:371–8.

48. Thomas DD, Navab M, Haake DA, Fogelman AM, Miller JN, Lovett MA. 1988. *Treponema pallidum* invades intercellular junctions of endothelial cell monolayers. Proc Natl Acad Sci U S A 85:3608–12.

49. Fitzgerald TJ, Johnson RC, Miller JN, Sykes JA. 1977. Characterization of the attachment of *Treponema pallidum* (Nichols strain) to cultured mammalian cells and the potential relationship of attachment to pathogenicity. Infect Immun 18:467–78.

50. Fitzgerald TJ, Johnson RC, Sykes JA, Miller JN. 1977. Interaction of *Treponema pallidum* (Nichols strain) with cultured mammalian cells: effects of oxygen, reducing agents, serum supplements, and different cell types. Infect Immun 15:444–52.

51. Fitzgerald TJ, Miller JN, Sykes JA. 1975. *Treponema pallidum* (Nichols strain) in tissue cultures: cellular attachment, entry, and survival. Infect Immun 11:1133–40.

52. Fitzgerald TJ, Cleveland P, Johnson RC, Miller JN, Sykes JA. 1977. Scanning electron microscopy of *Treponema pallidum* (Nichols strain) attached to cultured mammalian cells. J Bacteriol 130:1333–44.

53. Sykes JA, Miller JN, Kalan AJ. 1974. *Treponema pallidum* within cells of a primary chancre from a human female. Br J Vener Dis 50:40–4.

54. Sykes JA, Miller JN. 1971. Intracellular location of *Treponema pallidum* (Nichols strain) in the rabbit testis. Infect Immun 4:307–14.

55. Penn CW. 1981. Avoidance of host defences by *Treponema pallidum in situ* and on extraction from infected rabbit testes. J Gen Microbiol 126:69–75.

56. Lithgow KV, Church B, Gomez A, Tsao E, Houston S, Swayne LA, Cameron CE. 2020. Identification of the neuroinvasive pathogen host target, LamR, as an endothelial receptor for the Treponema pallidum adhesin Tp0751. mSphere 5:e00195–20.

57. Riley BS, Oppenheimer-Marks N, Radolf JD, Norgard MV. 1994. Virulent *Treponema pallidum* promotes adhesion of leukocytes to human vascular endothelial cells. Infect Immun 62:4622–5.

58. Azghani AO, Idell S, Bains M, Hancock RE. 2002. *Pseudomonas aeruginosa* outer membrane protein F is an adhesin in bacterial binding to lung epithelial cells in culture. Microb Pathog 33:109–14.

59. Fairman JW, Dautin N, Wojtowicz D, Liu W, Noinaj N, Barnard TJ, Udho E, Przytycka TM, Cherezov V, Buchanan SK. 2012. Crystal structures of the outer membrane domain of intimin and invasin from enterohemorrhagic *E. coli* and enteropathogenic *Y. pseudotuberculosis*. Structure 20:1233–43.

60. Krishnan S, Prasadarao NV. 2012. Outer membrane protein A and OprF: versatile roles in Gram-negative bacterial infections. FEBS J 279:919–31.

61. Oleastro M, Menard A. 2013. The role of *Helicobacter pylori* outer membrane proteins in adherence and pathogenesis. Biology (Basel) 2:1110–34.

62. Marra CM, Tantalo LC, Sahi SK, Dunaway SB, Lukehart SA. 2016. Reduced *Treponema pallidum*-specific opsonic antibody activity in hiv-infected patients with syphilis. J Infect Dis 213:1348–54.

63. Cruz AR, Ramirez LG, Zuluaga AV, Pillay A, Abreu C, Valencia CA, La Vake C, Cervantes JL, Dunham-Ems S, Cartun R, Mavilio D, Radolf JD, Salazar JC. 2012. Immune evasion and recognition of the syphilis spirochete in blood and skin of secondary syphilis patients: two immunologically distinct compartments. PLoS Negl Trop Dis 6:e1717.

64. Hawley KL, Cruz AR, Benjamin SJ, La Vake CJ, Cervantes JL, LeDoyt M, Ramirez LG, Mandich D, Fiel-Gan M, Caimano MJ, Radolf JD, Salazar JC. 2017. IFNgamma enhances CD64-potentiated phagocytosis of *Treponema pallidum* opsonized with human syphilitic serum by human macrophages. Front Immunol 8:1227.

65. Cox DL, Luthra A, Dunham-Ems S, Desrosiers DC, Salazar JC, Caimano MJ, Radolf JD. 2010. Surface immunolabeling and consensus computational framework to identify candidate rare outer membrane proteins of *Treponema pallidum*. Infect Immun 78:5178–94.

66. Avila-Nieto C, Pedreno-Lopez N, Mitja O, Clotet B, Blanco J, Carrillo J. 2023. Syphilis vaccine: challenges, controversies and opportunities. Front Immunol 14:1126170.

67. Lukehart SA, Shaffer JM, Baker-Zander SA. 1992. A subpopulation of *Treponema pallidum* is resistant to phagocytosis: possible mechanism of persistence. J Infect Dis 166:1449–53.

68. Salazar JC, Hazlett KR, Radolf JD. 2002. The immune response to infection with *Treponema pallidum*, the stealth pathogen. Microbes Infect 4:1133–40.

69. Moore MW, Cruz AR, LaVake CJ, Marzo AL, Eggers CH, Salazar JC, Radolf JD. 2007. Phagocytosis of *Borrelia burgdorferi* and *Treponema pallidum* potentiates innate immune activation and induces gamma interferon production. Infect Immun 75:2046–62.

70. White P, Haysom SF, Iadanza MG, Higgins AJ, Machin JM, Whitehouse JM, Horne JE, Schiffrin B, Carpenter-Platt C, Calabrese AN, Storek KM, Rutherford ST, Brockwell DJ, Ranson NA, Radford SE. 2021. The role of membrane destabilisation and protein dynamics in BAM catalysed OMP folding. Nature Communications 12:4174.

71. Weiser RS, Erickson D, Perine PL, Pearsall NN. 1976. Immunity to syphilis: passive transfer in rabbits using serial doses of immune serum. Infect Immun 13:1402–7.

72. Perine PL, Weiser RS, Klebanoff SJ. 1973. Immunity to syphilis. I. Passive transfer in rabbits with hyperimmune serum. Infect Immun 8:787–90.

73. Turner TB, Hollander DH. 1957. Biology of treponematoses. World Health Organization, Geneva, Switzerland.

74. Salazar JC, Vargas-Cely F, Garcia-Luna JA, Ramirez LG, Bettin EB, Romero-Rosas N, Amortegui MF, Silva S, Oviedo O, Vigil J, La Vake CJ, Galindo X, Ramirez JD, Martinez- Valencia AJ, Caimano MJ, Hennelly CM, Aghakhanian F, Moody MA, Sena AC, Parr JB, Hawley KL, Lopez-Medina E, Radolf JD. 2024. *Treponema pallidum* genetic diversity and its implications for targeted vaccine development: A cross-sectional study of early syphilis cases in Southwestern Colombia. PLoS One 19:e0307600.

75. Sena AC, Matoga MM, Yang L, Lopez-Medina E, Aghakhanian F, Chen JS, Bettin EB, Caimano MJ, Chen W, Garcia-Luna JA, Hennelly CM, Jere E, Jiang Y, Juliano JJ, Pospisilova P, Ramirez L, Smajs D, Tucker JD, Vargas Cely F, Zheng H, Hoffman IF, Yang B, Moody MA, Hawley KL, Salazar JC, Radolf JD, Parr JB. 2024. Clinical and genomic diversity of *Treponema pallidum* subspecies *pallidum* to inform vaccine research: an international, molecular epidemiology study. Lancet Microbe:100871.

76. Grillova L, Oppelt J, Mikalova L, Novakova M, Giacani L, Niesnerova A, Noda AA, Mechaly AE, Pospisilova P, Cejkova D, Grange PA, Dupin N, Strnadel R, Chen M, Denham I, Arora N, Picardeau M, Weston C, Forsyth RA, Smajs D. 2019. Directly sequenced genomes of contemporary strains of syphilis reveal recombination-driven diversity in genes encoding predicted surface-exposed antigens. Front Microbiol 10:1691.

77. Lieberman NAP, Lin MJ, Xie H, Shrestha L, Nguyen T, Huang ML, Haynes AM, Romeis E, Wang QQ, Zhang RL, Kou CX, Ciccarese G, Dal Conte I, Cusini M, Drago F, Nakayama SI, Lee K, Ohnishi M, Konda KA, Vargas SK, Eguiluz M, Caceres CF, Klausner JD, Mitja O, Rompalo A, Mulcahy F, Hook EW, 3rd, Lukehart SA, Casto AM, Roychoudhury P, DiMaio F, Giacani L, Greninger AL. 2021. *Treponema pallidum* genome sequencing from six continents reveals variability in vaccine candidate genes and dominance of Nichols clade strains in Madagascar. PLoS Negl Trop Dis 15:e0010063.

78. Molini B, Fernandez MC, Godornes C, Vorobieva A, Lukehart SA, Giacani L. 2022. B-cell epitope mapping of TprC and TprD variants of *Treponema pallidum* subspecies informs vaccine development for human treponematoses. Front Immunol 13:862491.

79. Silver AC, Dunne DW, Zeiss CJ, Bockenstedt LK, Radolf JD, Salazar JC, Fikrig E. 2013. MyD88 deficiency markedly worsens tissue inflammation and bacterial clearance in mice infected with *Treponema pallidum*, the agent of syphilis. PLoS One 8:e71388.

80. Murata T, Ohnishi M, Ara T, Kaneko J, Han CG, Li YF, Takashima K, Nojima H, Nakayama K, Kaji A, Kamio Y, Miki T, Mori H, Ohtsubo E, Terawaki Y, Hayashi T. 2002. Complete nucleotide sequence of plasmid Rts1: implications for evolution of large plasmid genomes. J Bacteriol 184:3194–202.

81. Parales J, Jr., Greenberg EP. 1993. Analysis of the *Spirochaeta aurantia flaA* gene and transcript. FEMS Microbiol Lett 106:245–51.

82. Beale MA, Marks M, Sahi SK, Tantalo LC, Nori AV, French P, Lukehart SA, Marra CM, Thomson NR. 2019. Genomic epidemiology of syphilis reveals independent emergence of macrolide resistance across multiple circulating lineages. Nat Commun 10:3255.

83. Chen W, Smajs D, Hu Y, Ke W, Pospisilova P, Hawley KL, Caimano MJ, Radolf JD, Sena A, Tucker JD, Yang B, Juliano JJ, Zheng H, Parr JB. 2021. Analysis of *Treponema pallidum* strains from China using improved methods for whole-genome sequencing from primary syphilis chancres. J Infect Dis 223:848–853.

84. Bioinformatics B. 2010. FastQC, *on* Babraham Institute. Accessed

85. Martin M. 2011. Cutadapt removes adapter sequences from high-throughput sequencing reads. EMBnetjournal 17:10–12.

86. Fraser CM, Norris SJ, Weinstock GM, White O, Sutton GG, Dodson R, Gwinn M, Hickey EK, Clayton R, Ketchum KA, Sodergren E, Hardham JM, McLeod MP, Salzberg S, Peterson J, Khalak H, Richardson D, Howell JK, Chidambaram M, Utterback T, McDonald L, Artiach P, Bowman C, Cotton MD, Fujii C, Garland S, Hatch B, Horst K, Roberts K, Sandusky M, Weidman J, Smith HO, Venter JC. 1998. Complete genome sequence of *Treponema pallidum*, the syphilis spirochete. Science 281:375–88.

87. Li H. 2018. Minimap2: pairwise alignment for nucleotide sequences. Bioinformatics 34:3094–3100.

88. LaFond RE, Centurion-Lara A, Godornes C, Rompalo AM, Van Voorhis WC, Lukehart SA. 2003. Sequence diversity of *Treponema pallidum* subsp. *pallidum tprK* in human syphilis lesions and rabbit-propagated isolates. J Bacteriol 185:6262–8.

89. LaFond RE, Centurion-Lara A, Godornes C, Van Voorhis WC, Lukehart SA. 2006. TprK sequence diversity accumulates during infection of rabbits with *Treponema pallidum* subsp. *pallidum* Nichols strain. Infect Immun 74:1896–906.

90. Lin MJ, Haynes AM, Addetia A, Lieberman NAP, Phung Q, Xie H, Nguyen TV, Molini BJ, Lukehart SA, Giacani L, Greninger AL. 2021. Longitudinal TprK profiling of *in vivo* and *in vitro*-propagated *Treponema pallidum* subsp. *pallidum* reveals accumulation of antigenic variants in absence of immune pressure. PLoS Negl Trop Dis 15:e0009753.

91. Danecek P, Bonfield JK, Liddle J, Marshall J, Ohan V, Pollard MO, Whitwham A, Keane T, McCarthy SA, Davies RM, Li H. 2021. Twelve years of SAMtools and BCFtools. Gigascience 10.

92. Cameron DL, Baber J, Shale C, Valle-Inclan JE, Besselink N, van Hoeck A, Janssen R, Cuppen E, Priestley P, Papenfuss AT. 2021. GRIDSS2: comprehensive characterisation of somatic structural variation using single breakend variants and structural variant phasing. Genome Biol 22:202.

93. 93. Team RC. 2021. R: A language and environment for statistical computing, *on* R Foundation for Statistical Computing. www.R-project.org. Accessed

94. Wickham H. 2016. ggplot2: Elegant graphics for data analysis. Springer-Verlag, New York.

95. Kenworthy AK, Schmieder SS, Raghunathan K, Tiwari A, Wang T, Kelly CV, Lencer WI. 2021. Cholera toxin as a probe for membrane biology. Toxins (Basel) 13.

